# Replication timing analysis in polyploid cells reveals Rif1 uses multiple mechanisms to promote underreplication in Drosophila

**DOI:** 10.1101/2021.04.19.440528

**Authors:** Souradip Das, Madison Caballero, Tatyana Kolesnikova, Igor Zhimulev, Amnon Koren, Jared Nordman

## Abstract

Regulation of DNA replication and copy number are necessary to promote genome stability and maintain cell and tissue function. DNA replication is regulated temporally in a process known as replication timing (RT). Rif1 is key regulator of RT and has a critical function in copy number control in polyploid cells. In a previous study (Munden et al., 2018), we demonstrated that Rif1 functions with SUUR to inhibit replication fork progression and promote underreplication (UR) of specific genomic regions. How Rif1-dependent control of RT factors into its ability to promote UR is unknown. By applying a computational approach to measure RT in Drosophila polyploid cells, we show that SUUR and Rif1 have differential roles in controlling UR and RT. Our findings reveal that Rif1 functions both upstream and downstream of SUUR to promote UR. Our work provides new mechanistic insight into the process of UR and its links to RT.

## Introduction

Replication of the genome is a highly regulated process that requires duplicating billions of bases of DNA with a high degree of accuracy. Failure to properly replicate genetic and epigenetic information each and every cell cycle can result in cell lethality or disease (Jackson et al., 2014). Regulation of DNA replication occurs largely at the initiation stage, where in late M and G1 phases of the cell cycle, the Origin Recognition Complex (ORC) facilitates loading of the MCM2-7 replicative helicase in an inactive state at all potential initiation sites (Bell & Labib, 2016). In S phase, a subset of these helicase complexes will be activated by Dbf4-Dependent Kinase (DDK). S-CDK activity then facilitates replisome assembly and the formation of bidirectional replication forks emanating from the replication start site (Siddiqui et al., 2013). To ensure genome stability, not all helicases are activated simultaneously during S phase (Collart et al., 2013; Mantiero et al., 2011). Rather, helicase activation is regulated temporally in a process known as Replication Timing (RT) (Gilbert, 2002; Rhind & Gilbert, 2013). RT refers to the precise time in S-phase when a given genomic region gets duplicated. RT is correlated with chromatin structure and activity: regions of the genome that replicate early tend to be accessible and transcriptionally active, whereas regions that replicate late tend to be less accessible and less transcriptionally active (Gilbert, 2002; Rhind & Gilbert, 2013).

RT is not merely a passive reflection of the chromatin state, but rather an actively regulated process. One major regulator of RT is the trans-acting factor Rif1 (<u>R</u>ap1-interacting <u>f</u>actor <u>1</u>). Rif1 controls genome-wide RT from yeast to humans (Armstrong et al., 2020; Cornacchia et al., 2012; Hayano et al., 2012; Peace et al., 2014; Seller & O’Farrell, 2018; Sreesankar et al., 2015; Yamazaki et al., 2012). The prevalent model for how Rif1 controls the RT program is based upon its conserved Protein Phosphatase 1 (PP1)-interaction motif. In this model, Rif1 recruits PP1 to loaded MCMs to oppose DDK-mediated helicase activation and to promote late replication (Davé et al., 2014; Hiraga et al., 2017; Hiraga et al., 2014). How Rif1 targets specific genomic regions or helicase molecules is unknown.

Despite the tight regulation of the DNA replication and RT programs, cell-type-specific regulation is necessary to accommodate cell-type-specific needs throughout development. For example, many cells of developing organisms are polyploid, having multiple copies of the genome in a single cell (Edgar & Orr-Weaver, 2001; Lilly & Duronio, 2005; Zielke et al., 2013). The genomes of polypoid cells, however, are not always fully replicated (Edgar & Orr-Weaver, 2001; Hua & Orr-Weaver, 2017; Nordman & Orr-Weaver, 2012; Spradling & Orr-Weaver, 1987). In Drosophila, most of the pericentric heterochromatin (PH) and certain euchromatic regions are underreplicated in polyploid cells. These underreplicated regions resemble chromosomal fragile sites found in mammals in that they lack replication origins, are late replicating, display tissue specificity and are associated with DNA damage (Andreyeva et al., 2008; Nordman et al., 2011; Nordman et al., 2014; Sher et al., 2012; Yarosh & Spradling, 2014). Underreplication is an actively regulated process, and in Drosophila the SUUR (<u>Su</u>ppressor of <u>U</u>nder<u>r</u>eplication) protein is required to promote UR (Belyaeva et al., 1998). SUUR is a potent inhibitor of replication fork progression and is promotes UR by inhibiting fork progression within specific regions of the genome (Nordman et al., 2014; Sher et al., 2012). SUUR, however, is unable to inhibit fork progression or promote UR on its own. Recently, we have shown that SUUR associates with Rif1 and recruits Rif1 to replication forks and that UR is completely dependent on Rif1 (Munden et al., 2018). Based on these data, we proposed that Rif1 acts downstream of SUUR to promote UR. Rif1 is known to regulate RT in multiple Drosophila tissues (Armstrong et al., 2020). It is unclear what contribution Rif1-dependent RT has on the promotion of UR, if any.

Here we explore the relationship between UR and late replication and what role SUUR and Rif1 play in controlling UR and RT. To investigate this, we applied a sorting-independent computational method to profile RT in Drosophila polyploid cells genome wide, which has not been feasible with traditional methods. In addition to generating the first high resolution genome-wide RT profiles of salivary gland and fat body tissues, we discovered that SUUR and Rif1 have differential roles in controlling UR and RT. Whereas both SUUR and Rif1 are essential for UR, only Rif1 has a substantial effect on RT. Critically, our results show that Rif1 functions both upstream and downstream of SUUR to promote UR depending on chromatin context. Together, our findings provide new mechanistic insight into the process of UR and its links to RT.

## Results

### TIGER can be used to generate RT profiles from large polyploid cells

To understand the relationship between underreplication and RT on a genome-wide scale requires high-resolution RT and UR profiles. While current high-resolution UR profiles already exist, or can be generated by Illumina-based sequencing, methods to generate high-resolution RT profiles require FACS sorting of precise S-phase populations, which is not always technically feasible for large polyploid cells. To overcome this technical challenge, we have utilized TIGER (<u>T</u>iming <u>I</u>nferred from <u>G</u>enome <u>R</u>eplication), a sequence-coverage based computational method that can measure RT without the need for sorting (Koren et al., 2021). Briefly, variations in DNA copy number driven by a modest percentage of cells in S phase within a population can be used to generate RT profiles. Therefore, RT profiles can be generated from Illumina sequence reads of a given cell- or tissue type if ∼10% or greater of the cells in the population are in S phase (Koren et al., 2014, 2021). TIGER has been used in mammalian cells to generate high-resolution genome-wide RT profiles and it rivals, or out performs, standard FACS-based methods to measure RT (Ding et al., 2020; Hulke et al., 2020; Massey et al., 2019).

To determine if TIGER could be adapted for a polytene tissue, we dissected salivary glands from third-instar larvae prior to wandering, extracted genomic DNA and Illumina sequenced the genomic DNA (Figure 1A). It was necessary to use 3^rd^ instar larvae prior to the wandering stage to ensure that >10% of cells in the tissue were in S phase. The raw sequencing files were run through a TIGER pipeline adapted for Drosophila to generate RT profiles (See Materials and Methods). As seem in Figure 1B, TIGER was able to generate profiles with characteristic peaks and troughs of a typical RT profile.

**Figure 1.**
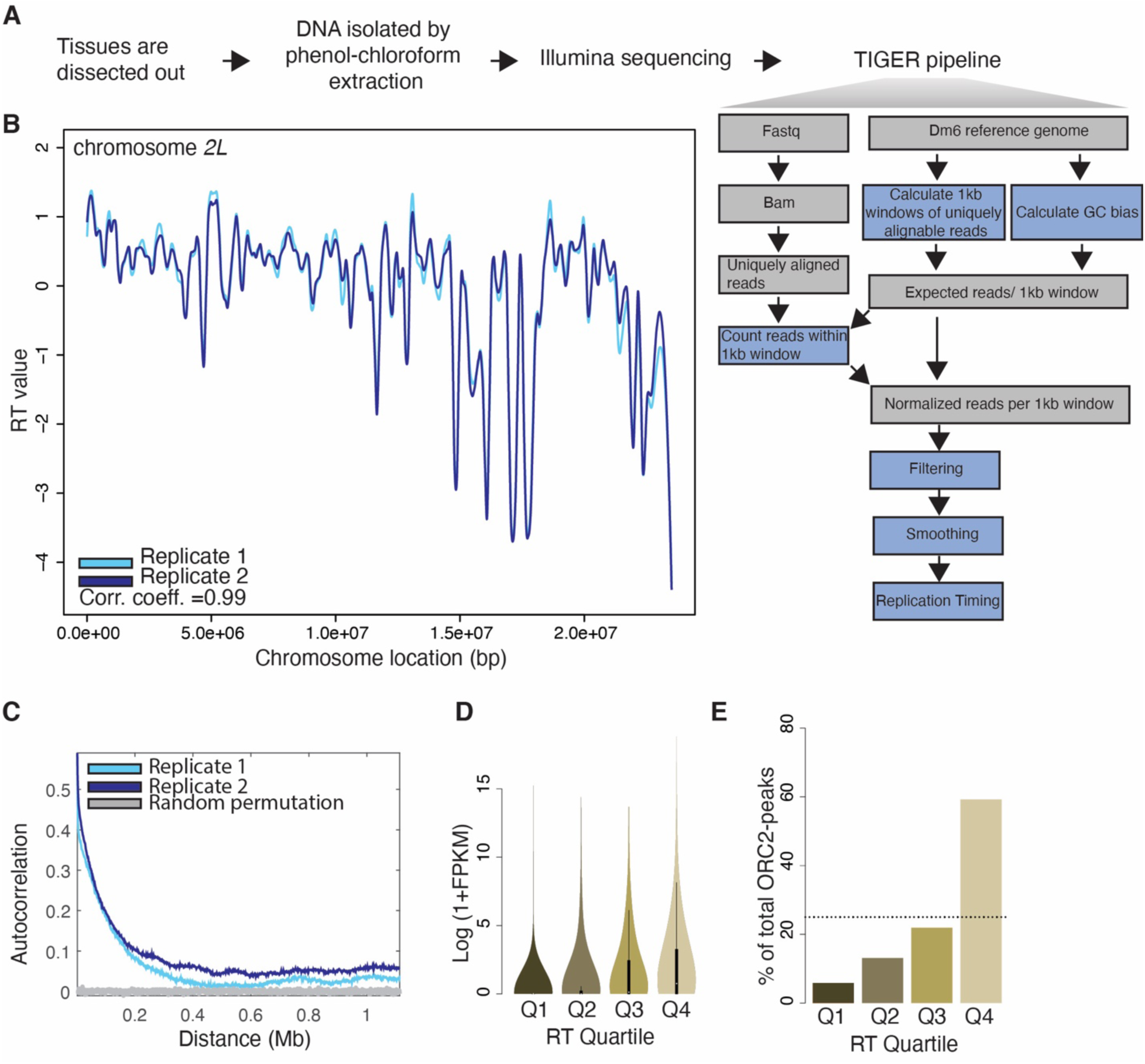
TIGER can generate RT profiles with high confidence from Drosophila polyploid tissues in a sorting-independent manner. A. Schematic outline of the workflow for preparing RT profiles through the TIGER pipeline: tissue dissection, DNA extraction, Illumina sequencing and TIGER pipeline. B. LOESS regression lines showing RT values for two biological replicates of wild-type salivary glands-replicate 1 (light blue) and replicate 2 (dark blue)-across the chromosome *2L* scaffold. See Figure S1A for all other chromosome arms. C. Autocorrelation values plotted for two replicates of wild-type salivary gland-replicate 1 (light blue) and replicate 2 (dark blue). Grey line depicts autocorrelation values for random permutation of RT values. D. RT windows from salivary gland were divided into quartiles with the lowest rawRT values in Q1 and the highest rawRT values in Q4. Average transcript FPKM values were calculated for every transcript within RT windows. The log2-transformed (1+FPKM) values were plotted for transcripts in each RT quartile in a violin plot. E. Bar graph showing the % of ORC2-peaks corresponding to RT windows grouped into RT quartiles in salivary gland. The dotted line marks the average % of the ORC2-peaks when equally distributed across quartiles.

Several lines of evidence indicate the profiles generated by TIGER truly represent RT in the larval salivary gland. First, LOESS-smoothed profiles of biological replicates were nearly superimposable with a correlation coefficient (r) = 0.99 (Figure 1B-C, Figure S1A-B). Second, replicates show a high degree of autocorrelation, which indicates a high degree of spatial continuity typical of replication timing data (Figure 1C). Third, transcripts emanating from genes in early replicating regions have a higher abundance than transcripts emanating from genes in late replicating regions (Figure 1D) (Nordman et al., 2011). Finally, early replicating regions are known to have a higher density of ORC2-binding sites (MacAlpine et al., 2010). To check if the relationship between ORC-distribution and RT holds true in our TIGER-generated salivary gland RT profiles, we used published ORC2 ChIP-seq data (Sher et al., 2012) and found that the percentage of ORC peaks is significantly higher in the earliest RT quartile (59.17%) compared to the latest RT quartile (5.81%) (Figure 1E). Taken together, the high autocorrelation patterning, positive correlation between early replication and high transcript abundance, positive correlation between early replication and ORC density establish TIGER as an effective solution for sorting-independent RT profiling of a polyploid tissue.

### RT profiles generated with TIGER correlate with RT profiles generated by the G1/S method

While the data presented in Figure 1 strongly suggests that the profiles generated by TIGER represent RT in the larval salivary gland, we wanted to compare these TIGER-generated profiles to RT profiles generated by conventional methods. Therefore, we compared directly TIGER-generated RT profiles in the larval salivary gland to RT profiles of larval wing discs and follicle cells of the adult ovary that our lab profiled using the G1/S method (Armstrong et al., 2020). If the TIGER-generated profiles truly reflect RT, then we would expect the RT profiles produced by TIGER to be comparable to these previously published. Comparing the profiles derived by two complementary methods required alternative normalization and smoothing steps (see Materials and Methods). Qualitatively, the RT profiles produced by both TIGER and G1/S method exhibit similar non-random patterning (Figure 2A-B, Figure S2A-B).

**Figure 2.**
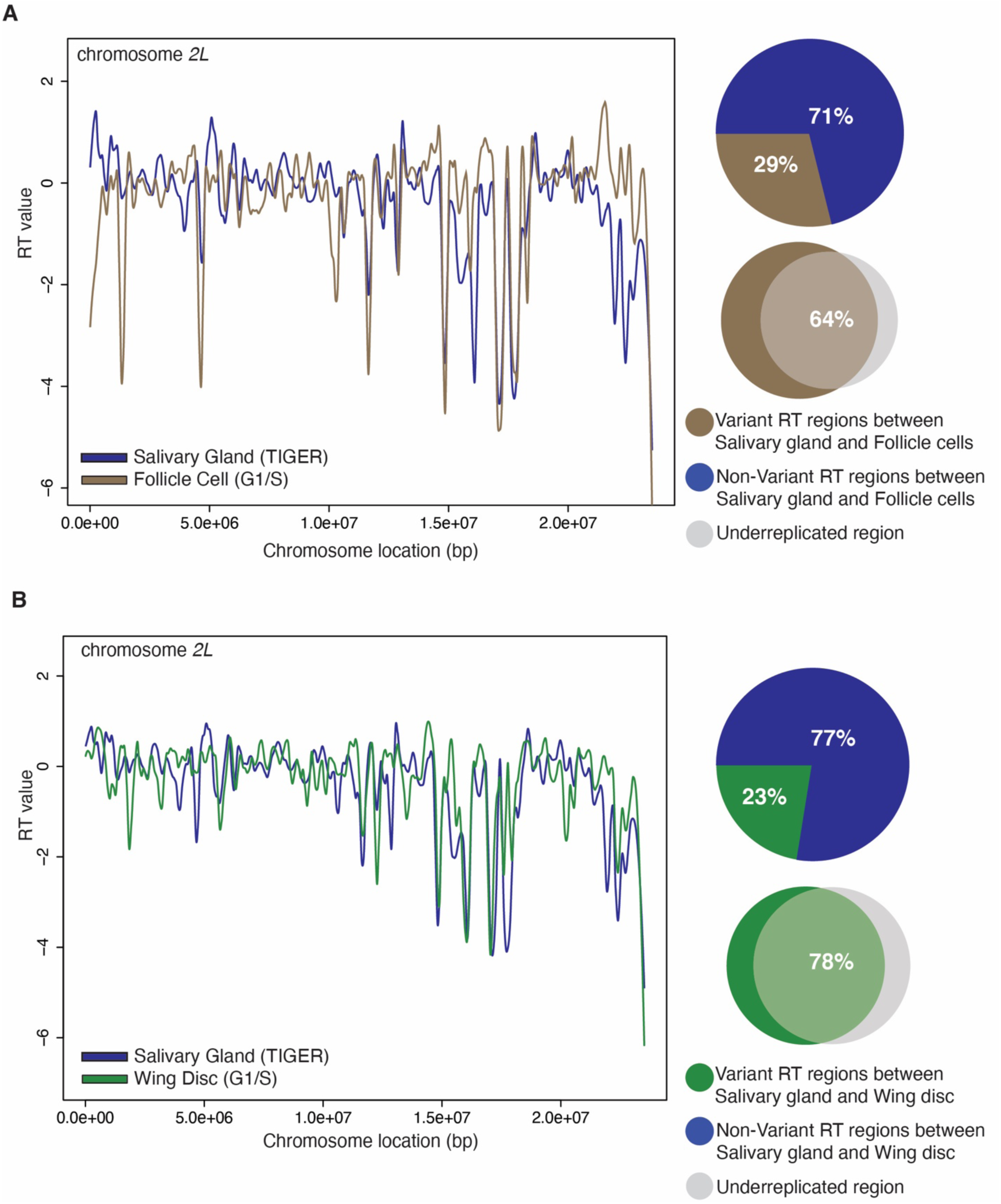
RT profiles generated with TIGER correlate with RT profiles generated by the G1/S method. A. LOESS regression lines showing average wild-type salivary gland (blue) and wild-type follicle cell (brown)-replication timing values across the chromosome *2L* scaffold. See Figure S2A for all other chromosome arms. Pie chart of all genomic windows of significantly different RT in follicle cell (brown) and unchanged RT in follicle cell (blue) relative to the salivary gland across the major chromosome scaffold. Venn diagrams comparing variant RT regions in the follicle cell (brown) relative to salivary gland, and underreplicated region from both euchromatin and pericentromeric heterochromatin region (grey). B. LOESS regression lines showing average wild-type salivary gland (blue) and wild-type wing disc (green)-replication timing values across the chromosome *2L* scaffold. See Figure S2B for all other chromosome arms. Pie chart of all genomic windows of significantly different RT in wing disc (green) and unchanged RT in wing disc (blue) relative to the salivary gland across the major chromosome scaffold. Venn diagrams comparing variant RT regions in the wing disc (green) relative to salivary gland, and underreplicated region from both euchromatin and pericentromeric heterochromatin region (grey).

Quantitatively we identified 71% of the genome in the follicle cells (Figure 2A) and 77% of the genome in the wing disc (Figure 2B) to exhibit similar RT relative to the salivary gland. Additionally, while characterizing the variant RT regions, we noted that majority of the variant regions, 64.45% and 78.28% respectively (Figure 2A-B), fall within known underreplicated regions of salivary glands. The differences in RT measured by TIGER and the G1/S method are consistent with expected differences due to cell-type specific changes in the RT program (Armstrong et al., 2020). Taken together, we conclude that TIGER is an effective method to generate genome-wide RT profiles from polyploid cells.

### Tissue-specific RT correlates with tissue-specific underreplication

Given that TIGER is an effective strategy to measure RT in polyploid tissues, it provides a tool to understand the relationship between tissue-specific RT and tissue-specific UR. Many larval tissues in Drosophila are polyploid and display tissue-specific underreplication (Nordman et al., 2011; Yarosh & Spradling, 2014). For example, the larval fat body reaches a ploidy of ∼256C with defined tissue-specific underreplicated sites. It is currently unknown, however, what the drivers of tissue-specific underreplication are. Therefore, we decided to profile larval fat body replication timing through TIGER to examine the relationship between RT and underreplication.

Larval fat body tissue was dissected from larvae 96h AEL (after egg laying). We used this time point to ensure that at least 10% of the cells in the tissue were in S phase (Hua et al., 2018). Genomic DNA was extracted, Illumina sequenced and RT profiles were generated using TIGER as described (see Materials and Methods). Consistent with our observation in salivary gland, biological replicates were highly correlated (Figure 3A, Figure S1A, S3A-B; Corr.

**Figure 3.**
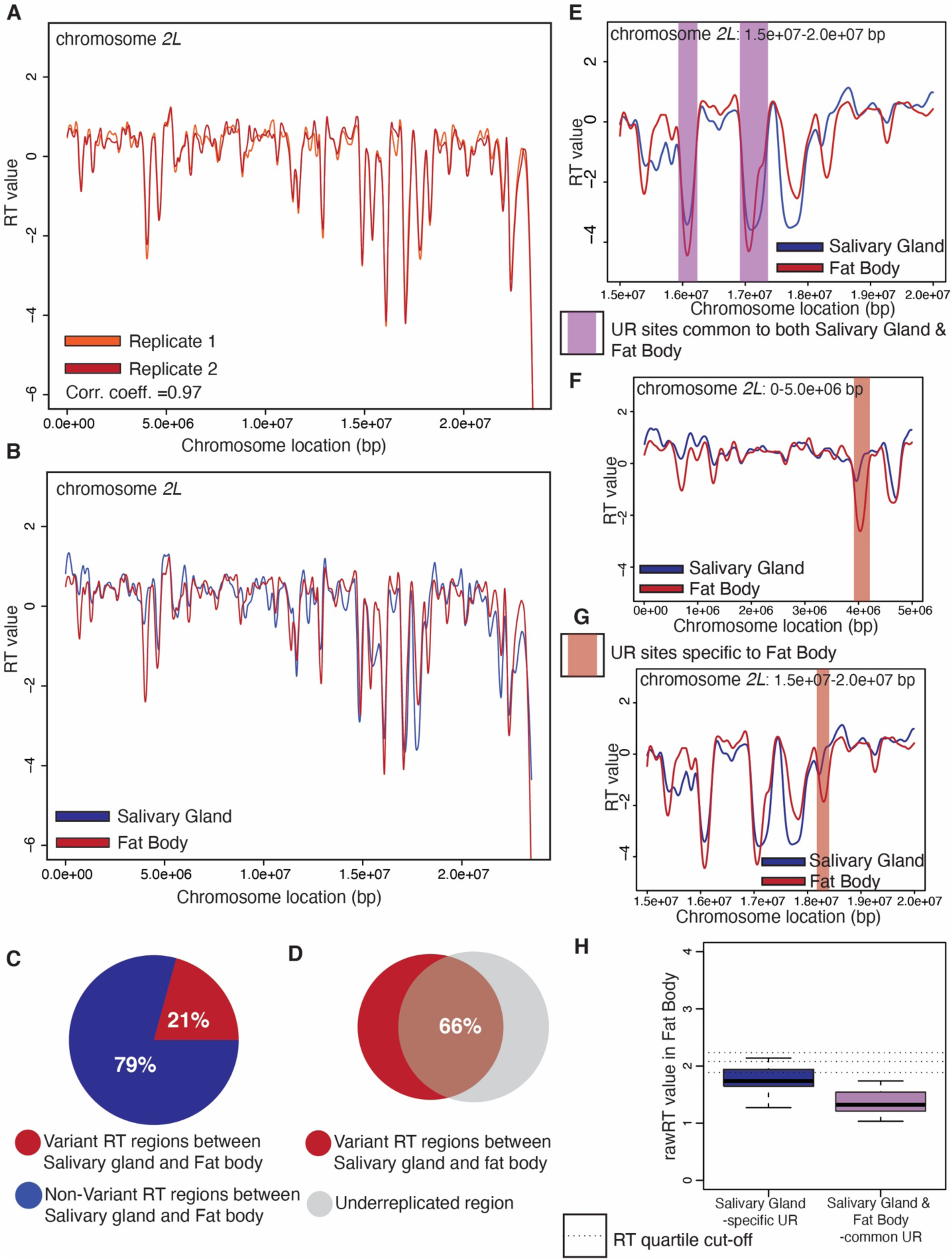
Tissue-specific RT correlates with tissue-specific underreplication. A. LOESS regression lines showing RT values for two biological replicates of wild-type larval fat body-replicate 1 (light red) and replicate 2 (dark red)-across the chromosome *2L* scaffold. See Figure S3A for all other chromosome arms. B. LOESS regression lines showing RT values for wild-type larval salivary gland (blue) and wild-type larval fat body (red)-across the chromosome *2L* scaffold. Each line represents the average of two biological replicates. See Figure S3B for all other chromosome arms. C. Pie chart of all genomic windows of significantly different RT in larval fat body (red) and unchanged RT in larval fat body (blue) relative to the salivary gland across the major chromosome scaffolds. D. Venn diagram comparing variant RT regions in the larval fat body (red) relative to salivary gland, and underreplicated regions from both euchromatin and pericentromeric heterochromatin (grey). E. LOESS regression lines showing RT values for wild-type salivary gland (blue) and wild-type larval fat body (red)-zoomed on two UR regions on chromosome *2L*, shared between salivary gland and larval fat body. The regions are highlighted with purple shading. F. LOESS regression lines showing RT values for wild-type salivary gland (blue) and wild-type larval fat body (red)-zoomed on an UR region on chromosome *2L*, specific to fat body. The region is highlighted with red shading. G. LOESS regression lines showing RT values for wild-type salivary gland (blue) and wild-type fat body (red)-zoomed on an UR region on chromosome *2L*, specific to fat body. The region is highlighted with red shading. H. Box plot quantifying the distribution of rawRT values of the UR regions in larval fat body. UR regions are either salivary gland specific (blue) or shared between salivary gland and fat body (purple). Dotted lines represent cut-off values for the RT quartiles.

Coefficient = 0.98) and early replicating regions in fat bodies have higher transcript abundance and increased ORC2 peak density relative to late replicating regions (Figure S3D-E). Previously published RNA-seq (Nordman et al., 2011) and ORC2 ChIP-seq (Hua et al., 2018) data from fat body was used for these analyses. The difference in transcript abundance between the earliest and latest RT quartile is not as pronounced in fat bodies as in salivary gland, however, this is consistent with previous observations demonstrating that transcripts in underreplicated regions (late replicating) of the fat body are substantially more expressed compared to that of salivary gland (Nordman et al., 2011).

Next, we quantified the variant RT regions between fat body and salivary glands and found that 21% of the genome showed RT differences between these two tissues (Figure 3B-C, Figure S3C). The majority of these variant regions (65.60%) fall within known underreplicated regions (Figure 3D). It is important to note that about 60% of these variants still remain in the similar RT quartile in both the tissues (Figure S3F). Importantly, we were able to identify tissue-specific RT variants that fall within underreplicated regions, raising the possibility that tissue-specific difference in RT could be correlated with tissue-specific UR (Figure 3C).

Underreplicated sites that reside within the euchromatic regions of the genome (eUR sites) fall into two categories: those that are tissue-specific and those that are common between multiple tissues (Nordman et al., 2011). Given the tissue-specific nature of underreplication, we reasoned that it provides an opportunity to determine if tissue-specific changes in RT correlate with a tissue-specific UR. To this end, we compared the TIGER-generated RT profiles of larval salivary gland and fat body tissues, which are known to contain both tissue-specific and common sites of UR. If tissue-specific UR correlates with tissue-specific RT, then we would expect to see significant changes in RT at these loci. As expected, UR sites that are shared between both tissues are late replicating in both tissues (Figure 3E). We next focused on the two sites that are underreplicated uniquely in the fat body tissue. These two fat body-specific UR sites replicate earlier in salivary gland compared to fat body, suggesting that tissue-specific UR is correlated with tissue-specific replication timing (Figure 3F-G).

Since there were only a limited number of fat body-specific UR sites, we also performed the inverse analysis, and compared the RT status of SG-specific UR sites in fat body. There are 40 UR sites present in SGs, of which 28 are specific to that tissue (Nordman et al., 2011).

We took the genomic coordinates these 40 UR sites and segregated them into two categories-salivary gland-specific URs and URs shared between salivary gland and fat body. Next, we extracted the RT value for each of these categories for fat body. If tissue-specific underreplication is correlated with tissue-specific RT, we would expect to see a later RT value (in the fat body) for sites that are common between the two tissues then sites that are specific for the salivary gland. The salivary gland-specific UR sites had replicated earlier in the fat body when compared to UR sites that are shared between both tissues (Figure 3H). This establishes a clear correlation between late replication and tissue-specific underreplication.

### SUUR and Rif1 have differential effects on RT and UR

Loss of SUUR or Rif1 function suppresses UR, however, the extent to which UR is suppressed in *SuUR* and *Rif1* mutants differs considerably. In an *SuUR* mutant, UR within the euchromatin regions is nearly completely suppressed while UR in the pericentric heterochromatin is only partially suppressed (Belyaeva et al., 1998; Munden et al., 2018). In contrast, UR within euchromatin and pericentromeric heterochromatin appears to be fully suppressed in a *Rif1* null mutant (Kolesnikova et al., 2020; Munden et al., 2018). To ask if these differences in UR could be caused by altered RT, we used TIGER to measure RT in *Rif1* and *SuUR* mutant larval salivary glands.

Genomic DNA was extracted from early third instar *SuUR* mutant salivary glands, sequenced and mapped reads were used to generate genome-wide RT profiles by TIGER. Similar to wild-type samples, biological replicates displayed high autocorrelation and high correlation between replicates, further supporting the robustness of TIGER (Figure 4A, Figure S1A, S4A, S4C). To quantitively measure the differences in RT between wild-type and *SuUR*, the genome was divided into non-overlapping ∼50kb windows and compared by ANOVA (see Materials and Methods). Comparison of RT profiles between wild-type and *SuUR* mutant salivary glands revealed that only 9% of the genome shows a significant difference in RT (Figure 4A). Given that RT profiles generated by TIGER rely on copy number measurements, we wanted to determine what fraction of RT changes reside within UR regions that are known to change copy number in an SUUR-dependent manner. 91.42% of the genomic regions that change RT in an SUUR-dependent manner fall within known UR regions and all but one region advances in RT in the *SuUR* mutant (Figure 4A). Critically, although there are substantial differences in copy number in these regions when comparing wild-type and *SuUR* mutant polytene chromosomes, there is only a minor change in RT within these regions (Figure S4E). Therefore, we conclude that SUUR has only a modest impact on RT and that TIGER can be used to accurately measure RT independently of large changes in copy number.

**Figure 4.**
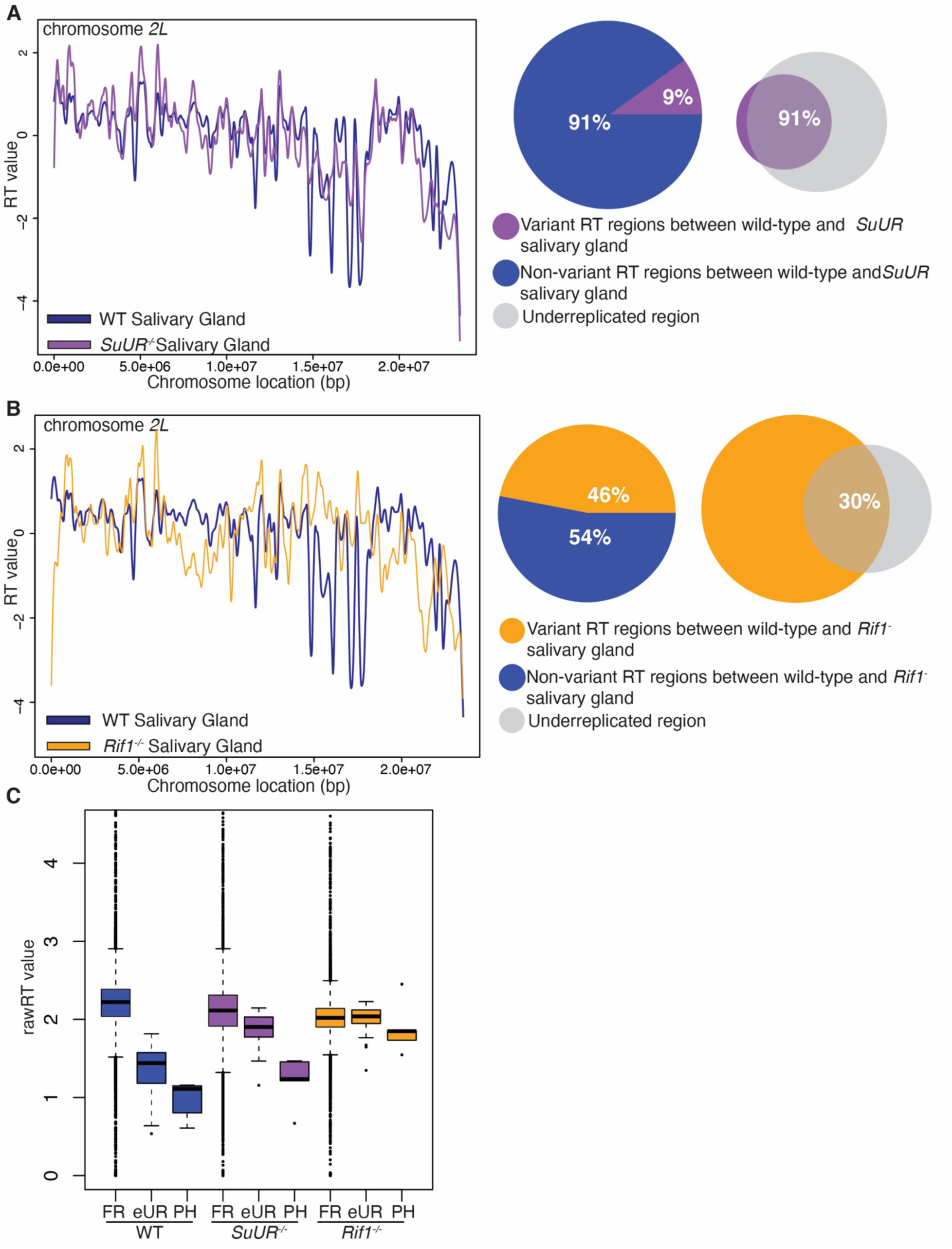
SUUR and Rif1 have differential effects on RT and UR. A. LOESS regression lines showing average wild-type (blue) and *SuUR* mutant (purple) salivary gland replication timing values across the chromosome *2L* scaffold. See Figure S4A for all other chromosome arms. Pie chart of all genomic windows of significantly different RT in an *SuUR* mutant (purple) and unchanged RT in an *SuUR* mutant (blue) relative to wild-type across the major chromosome scaffolds. Venn diagrams comparing variant RT regions in *SuUR* mutant (purple) relative to wild-type (purple) and underreplicated regions from both euchromatin and pericentromeric heterochromatin (grey) in salivary gland. B. LOESS regression lines showing average wild-type (blue) and *Rif1* mutant (yellow) salivary gland replication timing values across the chromosome *2L* scaffold. See Figure S3B for all other chromosome arms. Pie chart of all genomic windows of significantly different RT in a *Rif1* mutant (yellow) and unchanged RT in a *Rif1* mutant (blue) compared to wild-type, across the major chromosome scaffolds. Venn diagram comparing variant RT regions in *Rif1* mutant (yellow) relative to wild-type and underreplicated regions from both euchromatin and pericentromeric heterochromatin (grey) in salivary gland. C. Box plot quantifying the distribution of rawRT values of the genomic regions that are fully replicated (FR), underreplicated and comes from euchromatic region (eUR) or underreplicated pericentromeric heterochromatin (PH) in wild-type (blue), *SuUR* (purple) and *Rif1* mutant (yellow) salivary glands.

Next, we used TIGER to generate RT profiles from *Rif1* mutant larval salivary glands. Similar to wild-type and *SuUR* mutant data sets, highly correlated biological replicates displayed a high degree of autocorrelation (Figure 4B, Figure S4D, Figure S1A). Comparison of wild-type and *Rif1* mutant RT profiles of the larval salivary gland revealed that 46% of genomic regions had a significant change in RT (Figure 4B). In contrast to the *SuUR* mutant that had all but one region advance in RT, 32.60% of regions displayed advanced RT while 67.40% had delayed RT in the *Rif1* mutant. Of all the regions that displayed differential RT in the *Rif1* mutant relative to wild type, only 29.88% fell within known UR regions (Figure 4B). This indicates that, in contrast to SUUR, Rif1 functions as a global regulator of RT in the polyploid larval salivary gland similar to its function in other diploid cell types (Armstrong et al., 2020; Cornacchia et al., 2012; Foti et al., 2016; Hayano et al., 2012; Peace et al., 2014; Yamazaki et al., 2012).

To better understand the differences SUUR and Rif1 have on genome-wide RT, we generated histograms of RT values for wild-type, *SuUR* and *Rif1* mutant salivary glands (Figure S4F-H). While the distribution of RT values is nearly identical between wild-type and *SuUR* mutant salivary glands, *Rif1^-^* RT value distributions are significantly clustered toward the mean RT value, consistent with previous observations in mammalian cells (Cornacchia et al., 2012; Gnan et al., 2019; Hayano et al., 2012). Next, to determine the effect that SUUR and Rif1 have on RT, specifically within UR regions of the genome, we compared the RT values within UR regions. For this analysis we separated UR regions that fall within the euchromatic arms of the genome (eUR) from the underreplicated pericentric heterochromatin that is mappable by short-read sequencing (PH). As expected, the RT values for the UR regions were significantly later than the fully replicated regions of the genome in wild-type salivary glands, as UR regions are known to be late replicating (Figure 4C) (Nordman et al., 2011; Nordman & Orr-Weaver, 2012;

Nordman et al., 2014; Zhimulev et al., 1982, 2003). Similar to wild-type, eUR and PH regions of genome still displayed a pattern of late replication in the *SuUR* mutant (Figure 4C). While the difference in RT between the fully replicated and eUR or PH regions of the genome was not as substantial as in wild-type, this is likely due to changes in copy number within UR regions as UR regions in the *SuUR* mutant are still some of the latest replicating regions of the genome (Figure 4A). In stark contrast to wild-type and *SuUR* mutants, the RT values within the eUR regions were not significantly reduced relative to the fully replicated regions of the genome in the *Rif1* mutant (Figure 4C). The PH in the *Rif1* mutant, however, replicates significantly later than the fully replicated regions of the genome. Together these data indicate two key points. First, changes in copy number due to UR can be separated from changes in RT using TIGER. Second, SUUR and Rif1 have different effects on RT within UR regions. While eUR and PH are still late replicating in an *SuUR* mutant only PH is late replicating in the *Rif1* mutant. Therefore, we conclude that loss of SUUR function suppresses UR independently of RT, whereas loss of Rif1 function results in both loss of late RT and loss of UR only at eUR regions. This suggests that Rif1 functions upstream of SUUR at eUR regions to promote UR.

### Rif1 functions upstream of SUUR to promote underreplication of eUR regions

TIGER-generated RT profiles for wild type, *SuUR* and *Rif1* mutant salivary glands make two key predications that can be tested cytologically. First, Rif1 mutants should have a replication pattern that is clearly distinct from wild-type or *SuUR*-mutant salivary glands along the euchromatic arms. Second, *SuUR* binding to eUR sites (along the euchromatic arms of chromosomes) should largely be abolished since they are no longer late replicating in the *Rif1* mutant. SUUR binding to pericentric heterochromatin, however, should be largely unaffected in *Rif1* mutant polytene salivary glands. In support of the first prediction, the global pattern of EdU incorporation into chromosome arms of *Rif1* mutant salivary gland polytene chromosomes has recently been shown to differ significantly from that of wild-type or *SuUR* mutant polytene chromosomes. Sites that are normally late replicating and underreplicated in wild-type chromosomes appear to replicate earlier in S phase in a *Rif1* mutant (Kolesnikova et al., 2020). To extend this finding, we monitored the completion of replication of salivary glands by performing polytene squashes and staining chromosomes with anti-PCNA antibody to mark sites of active replication. PCNA labeling within the *35A* to *36D* region on chromosome *2L*, which contains several UR loci, was monitored for replication in different stages of S phase. In the *SuUR* mutant, UR regions at *35B*, *35E* and *36D* all complete replication in late S phase, consistent with our TIGER measurements of RT (Figure 4A). Additionally, the normally underreplicated region at *36C* completed replication prior to the underreplicated region at *36D* (Figure 4A). Consistent with this cytological observation, our TIGER RT profiles revealed that *36C* has an earlier RT value then *36D*. In contrast, these same regions complete replication in mid S phase in the *Rif1* mutant rather than late S phase in the *SuUR* mutant (Figure 5A). These specific replication timing differences validate the RT values we measured by TIGER.

**Figure 5.**
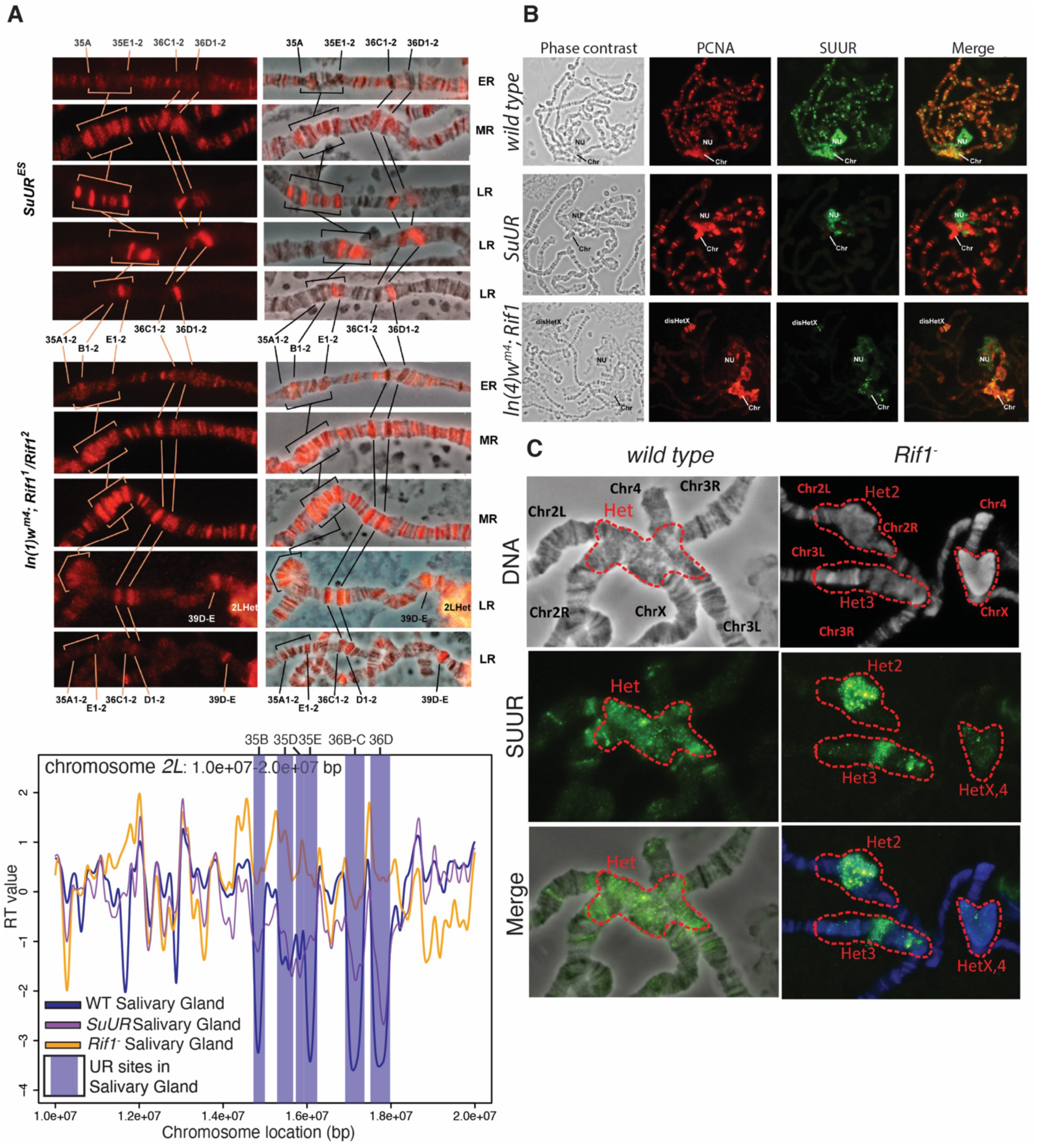
Rif1 functions upstream of SUUR to promote underreplication of euchromatic UR regions. A. PCNA staining patterns in the *35A-36D* region of *chr2L*. *35E, 36D and 36D* are normally underreplicated. TIGER-generated RT profiles of the same region are shown for reference. ER – early replicating, MR – mid S phase replication, LR – late replicating. B. anti-SUUR (green) and anti-PCNA (red) immunostaining of late replicating nuclei of the following genotypes: Oregon R (wt), *SuUR, In(1)wm4*; *Rif1*. Note that the *In(1)wm4* chromosome has an inversion that translocates a portion of heterochromatin on the X chromosome to *3C*. Chr – chromocenter, Chr3Het – Chromosome 3 heterochromatin, disHetX – distal heterochromatin of chromosome X transferred by inversion *In(1)wm4* to the region 3C, NU – nucleolus. C. anti-SUUR (green) and anti-PCNA (red) immunostaining of late replicating nuclei of the following genotypes: Oregon R (wt), *Rif1*. Chr – chromosome, Het – heterochromatin.

Since UR is abolished in a *Rif1* mutant and our RT profiles reveal that normally eUR regions replicate significantly earlier in a *Rif1* mutant (Figure 4B and 4C), this raised the possibility that Rif1 acts upstream of SUUR to promote UR of the eUR regions. Because SUUR is known to target late replicating regions of the genome, it should no longer bind to those regions if they replicate earlier in S phase. SUUR, however, should still bind to the late replicating pericentric heterochromatin (PH) in a *Rif1* mutant as SUUR was shown to localize to heterochromatin in a *Rif1* mutant and heterochromatin replicates late in *Rif1* mutant salivary glands (Munden et al., 2018). We monitored the localization of SUUR in wild-type and *Rif1* mutant polytene chromosomes using a SUUR antibody. As predicted, SUUR does not bind to the eUR regions in *Rif1* mutant salivary glands (Figure 5B). In contrast, SUUR still localized to the pericentric heterochromatin in *Rif1* mutant salivary gland chromosomes (Figure 5B and 5C). We did observe, however, that the pattern of SUUR localization to the chromocenters was slightly altered in a *Rif1* mutant (Figure 5C). SUUR was not evenly distributed throughout the entire chromocenter, rather some regions of the chromocenter were stained more brightly than others and SUUR was often clustered in puncta (Figure 5C). Together, these cytological data complement our TIGER-generated RT profiles and indicate that Rif1 can act upstream of SUUR to promote RT within the eUR regions of the genome.

## Discussion

To understand the relationship between UR and RT, we have applied TIGER, a sorting-independent computational method to profile RT genome wide, to successfully measure RT in several polyploid tissues. By comparing genome-wide RT profiles between wild type, *SuUR* and *Rif1* mutants, we were able to further our understanding of the mechanisms Rif1 employs to promote underreplication. We have found that SUUR and Rif1 differentially control RT, Furthermore, in addition to functioning downstream of SUUR to promote UR (Munden et al. 2018), Rif1 can also function upstream of SUUR to promote underreplication by promoting late replication of specific genomic regions. Additionally, our work has further emphasized the link between late replication and UR, demonstrating that late replication is essential for underreplication. Finally, we showed that tissue-specific UR is correlated with tissue-specific RT, which highlights how replication programs can be modulated during development to ensure proper tissue function.

Methods to measure the genome-wide patterns of RT are dependent on FACS to isolate S and G1 phase populations (Armstrong et al., 2018; Hulke et al., 2020; Ryba et al., 2011; Marchal et al., 2018). FACS of large polyploid cells for RT profiling, however, is not efficient and this limitation has prevented profiling polyploid cells of the larval salivary gland, which has served as an important model for genome biology. Through the utilization of TIGER, we have overcome this challenge. This sorting-independent protocol is the latest addition to the repertoire of RT profiling methods and has proven to be both time saving and cost effective. We were able to generate TIGER-based RT profiles of multiple polyploid cells and several lines of evidence indicate that these profiles reflect true genome-wide RT profiles. RT profiles are highly reproducible between biological replicates, have a high degree of autocorrelation as a function of chromosome position, are correlated with transcript abundance and frequency of ORC2 binding sites and are similar to RT profiles of diploid tissues generated by the G1/S method.

TIGER-generated RT profiling of larval salivary glands revealed that while SUUR and Rif1 are both necessary for UR, they have significantly different effects on RT. While SUUR is responsible for a significant portion of UR (Makunin et al., 2002; Munden et al., 2018; J. T. Nordman & Orr-Weaver, 2015), it has only a modest effect on RT. In contrast, nearly 50% of the genome is dependent on Rif1 for proper RT. The fraction of the salivary gland genome that is dependent on Rif1 for RT is significantly greater than what was observed for Drosophila follicle cells or wing discs (Armstrong et al., 2020). While differences in the statistical methods used to perform variant calling could be responsible for subset of these differences, it is clear that Rif1 differentially affects RT in these three cell types. This is consistent with our previous observations that Rif1 controls RT in a cell-type-specific manner (Armstrong et al., 2020). It is still unknown, however, how Rif1’s activity is regulated to establish cell-type-specific patterns of RT.

Our previous work revealed that SUUR recruits Rif1 to replication forks to inhibit fork progression (Munden et al., 2018). RT profiling of Rif1 mutant salivary glands and SUUR localization in *Rif1*-mutant salivary glands indicates that Rif1 functions upstream of SUUR to promote UR. Thus, we propose a revised model for Rif1-mediated UR in eUR and PH regions (Figure 6). First, Rif1 acts upstream of SUUR to promote late replication of specific eUR loci. Subsequently, SUUR localizes to a subset of those late replicating regions to inhibit replication fork progression in conjunction with Rif1. In the absence of Rif1, however, eUR regions that are normally underreplicated lose their late RT status and SUUR is unable to target replication forks within these regions. The PH UR sites, however, are still late replicating in absence of Rif1 likely due to additional factors that are dependent on H3K9 methylation status (Armstrong et al., 2019). SUUR still localizes to these PH regions, but is unable to inhibit fork progression and promote UR because Rif1 is not present and SUUR is unable to inhibit fork progression in the absence of Rif1 (Munden et al., 2018). This model is in agreement with multiple observations indicating that SUUR binding to euchromatic and pericentric heterochromatin is different and that the genetic requirements for UR in euchromatin and pericentric heterochromatin are unique (Armstrong et al., 2019; Guarner et al., 2017).

**Figure 6.**
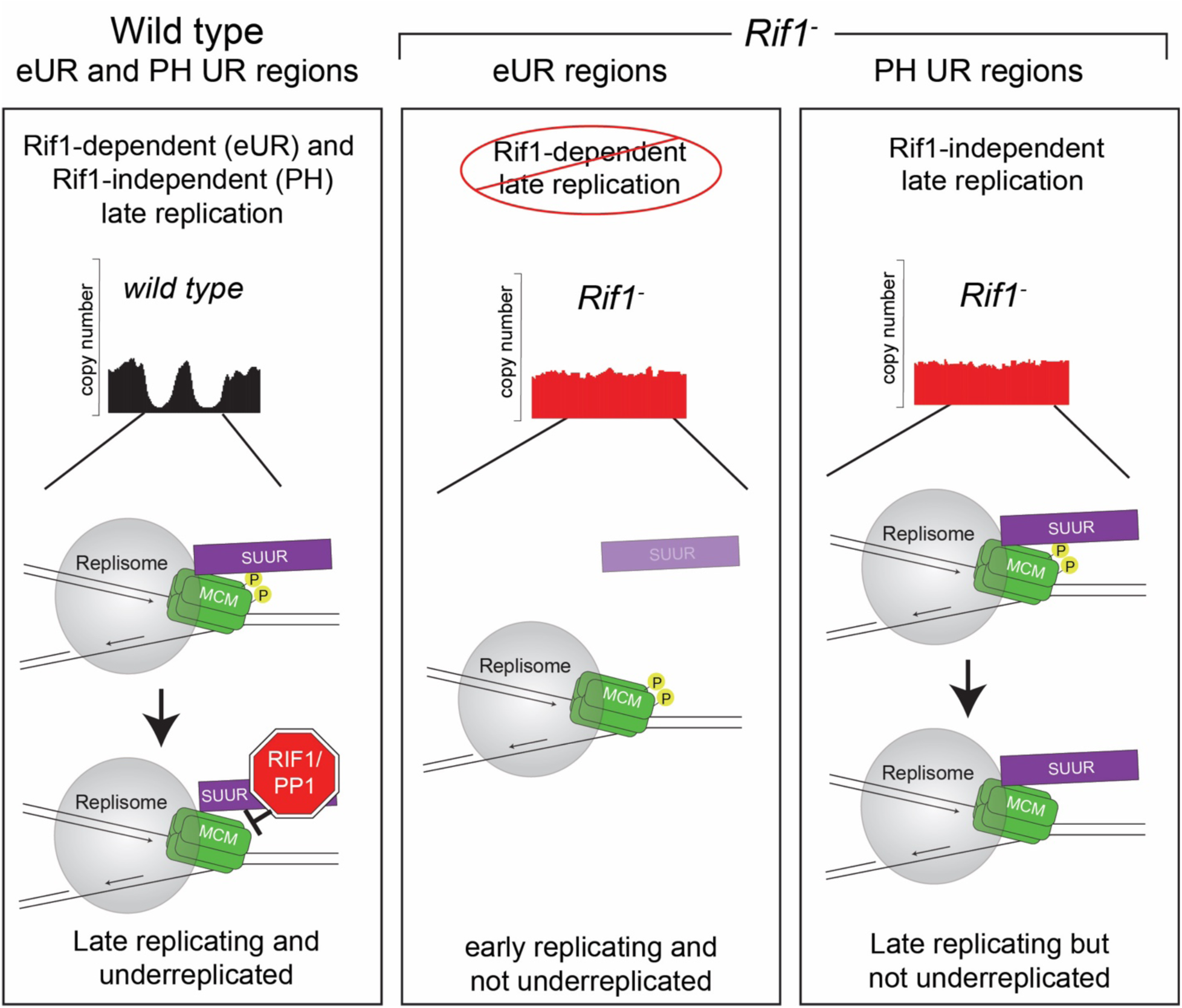
A model for Rif1-dependent promotion of underreplication. In this revised model, Rif1 first promotes late replication of certain genomic regions. Next, SUUR promotes underreplication in a subset of those late-replicating regions by recruiting Rif1/PP1 to replication forks to inhibit fork progression. Without Rif1, eUR regions replicate early and SUUR is unable to target replication forks in those regions. PH regions, however, remain late replicating in the absence of Rif1. Therefore, SUUR is able to target replication forks within PH regions. Without Rif1, SUUR cannot inhibit fork progression to promote UR.

It is still unknown how exactly SUUR targets specific late-replicating regions of the genome to promote UR. Additionally, it is unclear what aspects of chromatin structure and/or function change during development to generate tissue-specific patterns of UR. Data presented here demonstrate that tissue-specific RT correlates with tissue-specific UR. Additionally, our work further emphasizes the relationship between late replication and UR. While late replication appears to be necessary for UR, it is not sufficient. Therefore, in addition to replication timing there must be additional factors that are essential for SUUR-mediated UR.

## Materials and Methods

### Strains

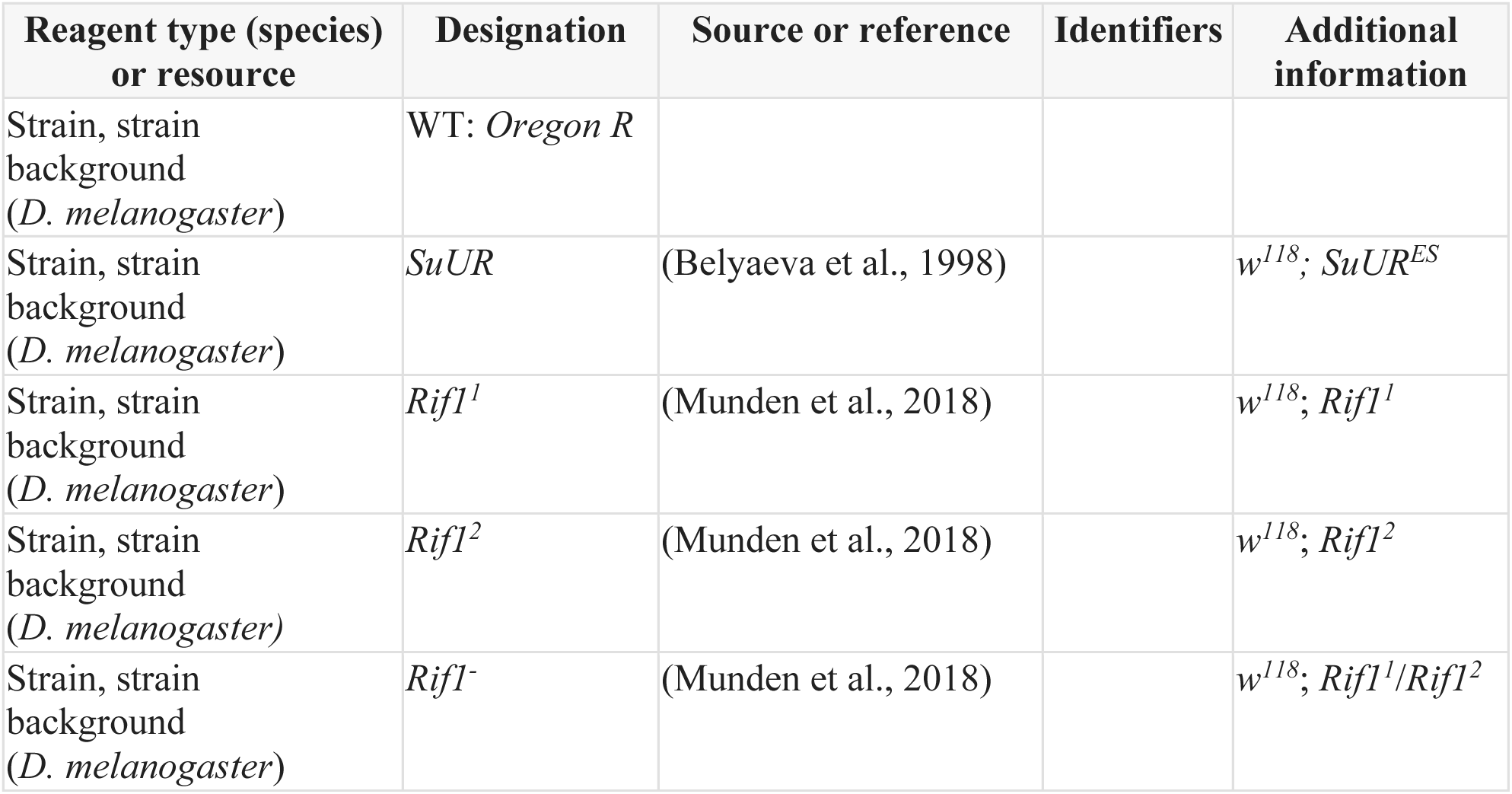

### Genomic DNA sequencing

Salivary glands were dissected in EBR from ∼20 3rd instar female larvae prior to wandering per genotype, wild type -*OregonR, Rif1 - Rif^1^1/Rif^2^,* and *SuUR* - *SuUR^ES^*. Fat bodies were dissected in EBR from 25 OregonR female larvae per replicate 96 hours after egg laying (AEL). Tissues were pelleted, resuspended in LB3, dounce homogenized and sonicated using a Bioruptor 300 (Diagenode) for 10 cycles of 30’’ on and 30’’ off at maximal power. Lysates were treated with RNase and Proteinase K and genomic DNA was isolated by phenol-chloroform extraction. Illumina libraries were prepared using NEBNext DNA Ultra II DNA Library Prep Kit for Illumina (New England Biolabs) following the manufacturers protocol. Barcoded libraries were sequenced using Novaseq 6000 paired-end 150bp sequencing.

### Bioinformatics

#### RT generation

Bam files were aligned to dm6 (Dos Santos et al., 2015) by BWA mem (v0.7.17) (Li, 2013) and duplicate reads are marked with Picard Tools command ‘MarkDuplicates’.

Coordinates of uniquely-mapping, non-duplicate, reads with a MAPQ >10 were extracted with samtools (Li et al., 2009) view (v1.11) (-F 1024 -F 256 -F 128 -q 10). All the salivary gland and fat body samples processed were in biological duplicates. RT values for wing disc and follicle cell were from previously published data (Armstrong et al., 2020).

RT was generated from a modified version of TIGER (Koren et al., 2021). Alignability filtering was performed against dm6 using a read length of 100bp. The uniquely aligning sample reads were then partitioned into windows of 1000 uniquely alignable base pairs. GC correction was performed with standard segmentation (TIGER command ‘TIGER_segment_filt’, using the MATLAB function ‘segment’) parameters which temporarily removed eUR and PH regions for determining GC content bias. GC correction was then applied to all data (PH and eUR included).

From the GC-corrected data in 1kbp windows, eUR regions were defined in WT fat body and salivary gland samples as regions of continuous low DNA copy number. In this, the raw DNA copy number data in 1kb windows were segmented (TIGER command ‘TIGER_segment_filt’, R2 = 0.04, standard deviation threshold = 1) to identify regions of at least one standard deviation below the mean. From these regions, only those ≥ 50 kbp in length without gaps ≥10 kbp (≥10 continuous 1kbp windows above the segmentation standard deviation cutoff) were called as eUR regions. This method provided the most accurate prediction of eUR regions by visual inspection. These predicted eUR regions overlapped with the smaller panel of previously published UR regions (Nordman et al., 2011) in fat body and salivary glands. The predicted and previously published eUR regions were merged with bedtools (Quinlan & Hall, 2010) merge (v2.29.2) to finalize eUR zones in fat body and salivary gland.

Standard TIGER data filtering removes outlier windows attributed to noise or copy number variations. To still filter outliers, eUR and PH regions must be removed and filtered separately from the rest of the chromosome (arm regions). The eUR (as defined in the previous paragraph) and PH regions were removed from the GC-corrected data. In the arm-only regions, outliers were removed with segmentation (standard deviation threshold = 2, R2 = 0.06). eUR regions then separately filtered for outliers in a similar manner (standard deviation threshold = 3, R2 = 0.06). All segmentation parameters were chosen to optimize outlier removal via visual interpretation.

The segmentation filtered arm, PH, and eUR regions were merged to form the final rawRT data. This data was used for all the statistical analysis including variant calling between salivary gland and fat body samples. RT values were generated by smoothing the filtered data with a cubic smoothing spline (MatLab command ‘csaps’, smoothing parameter = 1x10-15). Only zones of >20 continuous 1kbp windows were included and smoothing was not performed over gaps of >5kpb. The smoothed profiles were then normalized to an individual chromosomal mean of zero and a standard deviation of one.

In comparing TIGER profiles of salivary glands and fat body to G1/S profiles of wind disc and follicle cells, the variable copy number of eUR and PH regions distorts normalization. Therefore, a second normalization was performed based only on the arm regions of chromosomes. The arm regions were first normalized to a mean of zero and a standard deviation of one. The mean shift and standard deviation values based on the arm regions were then applied to the eUR and PH regions. All regions were then merged and smoothed for visualization using identical parameters.

#### Variant calling

The stats (v3.6.2) statistical package in R was used to identify 50-kb windows with significantly altered rawRT values, by one-way ANOVA test [aov, P value adjusted for multiple testing with Bonferonni post-HOC correction {adjusted P value < (0.01/no. of testing)}]. Adjacent windows were merged and regions >200kb in length were called as variant.

#### Correlation and autocorrelation

Correlation of RT profiles is generated from MATLAB command ‘corr’ (values are Pearson’s r). Autocorrelation was calculated using the MATLAB command ‘autocorr’ (number of lags = 1000).

#### RNA sequencing analysis

RNA-seq data for salivary gland was obtained from NCBI Gene Expression Omnibus (GEO) (http://www.ncbi.nlm.nih.gov/geo/) under the reference series GSE31900. RNA-seq data for fat body was obtained from NCBI Gene Expression Omnibus (GEO) (http://www.ncbi.nlm.nih.gov/geo/) under the reference series GSE25025. TopHat default parameters (v2.1.1) were used to align reads to the dm6 version of the Drosophila genome.

Transcriptomes were generated using Cufflinks (v2.2.1). Transcript FPKM values for each RT window were generated by calculating the mean FPKM values for all the transcript regions overlapping the window. Overlap was determined by BEDTools intersect (v2.27.1) with -f 0.5 parameters.

#### ORC2 ChIP-seq analysis

ORC2 peak data for salivary gland was obtained from NCBI Gene Expression Omnibus (GEO) (http://www.ncbi.nlm.nih.gov/geo/) under the reference series GSE31900. ORC2 peak data for fat body was obtained from NCBI Gene Expression Omnibus (GEO) (http://www.ncbi.nlm.nih.gov/geo/) under the reference series GSE90916. Coordinates were converted to dm6 coordinates using the UCSC liftOver tool (Karolchik et al., 2004).

#### Miscellaneous bioinformatics

BEDTools intersect (v2.27.1) was used to determine overlap of RT windows with −f 0.5 parameters. Merging of adjacent RT windows was done by BEDTools merge (v2.27.1) with -d 11 parameters. Quartile cut-offs for each genotype or tissue were calculated by the stats (v3.6.2) statistical package. Coordinates from dm3 were converted to dm6 using the UCSC liftOver tool (Karolchik et al., 2004).

#### Indirect Immunofluorescent Staining

For immunostaining of polytene chromosome squashes, salivary glands from wandering third instar larvae were dissected in PBST (137 mM NaCl, 3 mM KCl, 8 mM NaH2PO4 and 2 mM KH2PO4; 0.1 % Tween-20). Glands were then transferred into a formaldehyde-based fixative (0.1 M NaCl, 2 mM KCl, 10 mM NaH2PO4, 2 % NP-40, 2 % formaldehyde) for 1 min. Salivary glands were placed in an acetic acid–formaldehyde mix (45 % acetic acid, 3.2 % formaldehyde) for 1 min and squashed in 45 % acetic acid. Squashes were snap-frozen in liquid nitrogen and coverslips were removed. Slides were incubated in 70 % ethanol for 5 min twice and stored in 70 % ethanol at −20°С. Slides were first washed three times in PBST for 5 min. Primary antibodies were added in a blocking solution (0.1 % BSA in PBST) and incubated in humid chamber for 2 h at room temperature. The primary antibody dilutions used were as follows: rabbit polyclonal anti-SUUR (E-45) (Makunin et al. 2002), 1:50; mouse monoclonal anti-PCNA (PC10, Abcam, ab29). Then, squashes were washed in PBST and incubated in secondary antibody (Alexa Fluor 488-conjugated goat anti-rabbit and Alexa Fluor 568–conjugated goat anti--mouse IgG antibodies, 1:500; Thermo Fisher Scientific) in blocking solution for 1h. Squashes were mounted in VectaShield (Vector Laboratories) DAPI medium with 15 μg/mL DAPI. Images were acquired using an Olympus BX51 microscope equipped with a 100×/1.30 Uplan FI Ph3 oil objective and a DP70 camera.

### Data access

Data sets described in this manuscript can be found under the GEO accession number: GSE172375

## Acknowledgements

This work was supported by a National Science Foundation grant (MCB-818019 to J.T.N); National Institutes of Health (DP2-GM123495 to A.K.) and the National Science Foundation (MCB-1921341 to A.K.); and a Joint Russian–German grant from the Russian Foundation for Basic Research (No. 20-54-12016) and Deutsche Forschungsgemeinschaft Schu762/12-1. Illumina sequencing was performed at the VANTAGE core at Vanderbilt University Medical Center, which is supported by the CTSA Grant (5UL1 RR024975-03), the Vanderbilt-Ingram Cancer Center (P30 CA68485), the Vanderbilt Vision Center (P30 EY08126), and NIH/NCRR (G20 RR030956). We thank Terry Orr-Weaver, Robin Armstrong and members of the Nordman lab for comments on the manuscript.

## Competing interests

The authors express no conflicts of interest

## Supplementary Figures

**Figure S1.**
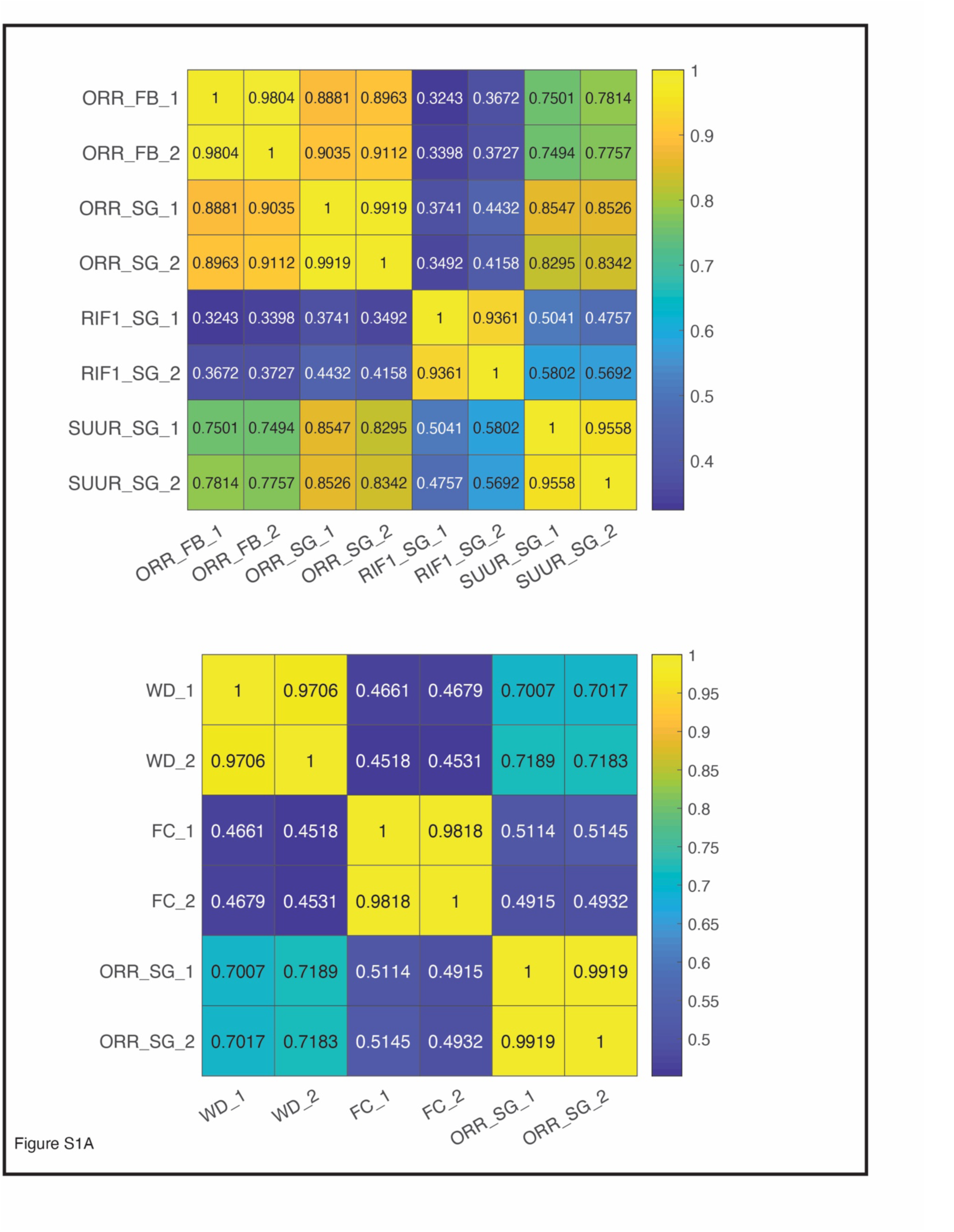
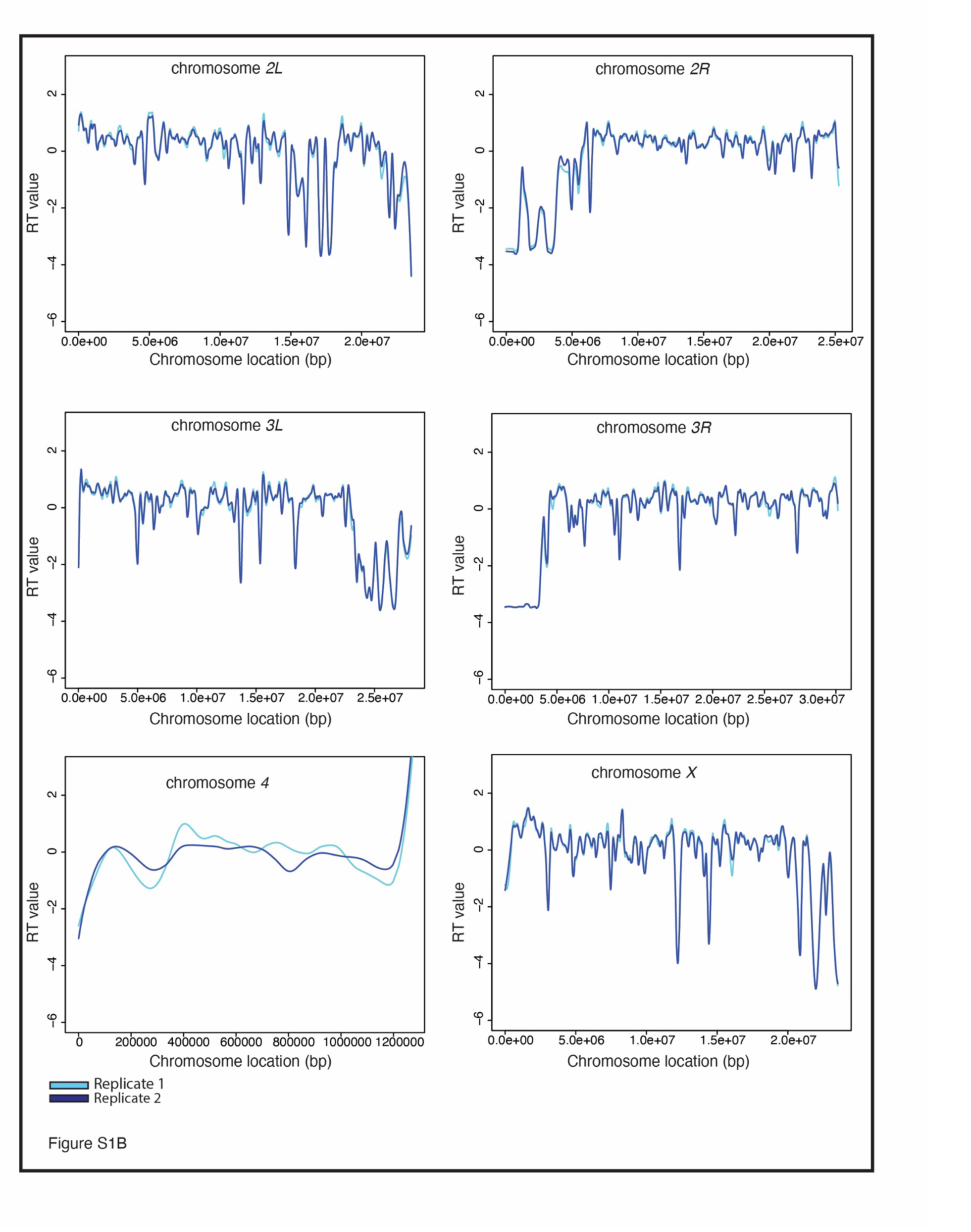
TIGER can generate RT profiles with high confidence from Drosophila polyploid tissues in a sorting-independent manner. A. Correlation matrix for all TIGER samples. B. LOESS regression lines showing RT values for two biological replicates of wild-type salivary glands replicate 1 (light blue) and replicate 2 (dark blue)- across all major chromosome scaffold.

**Figure S2.**
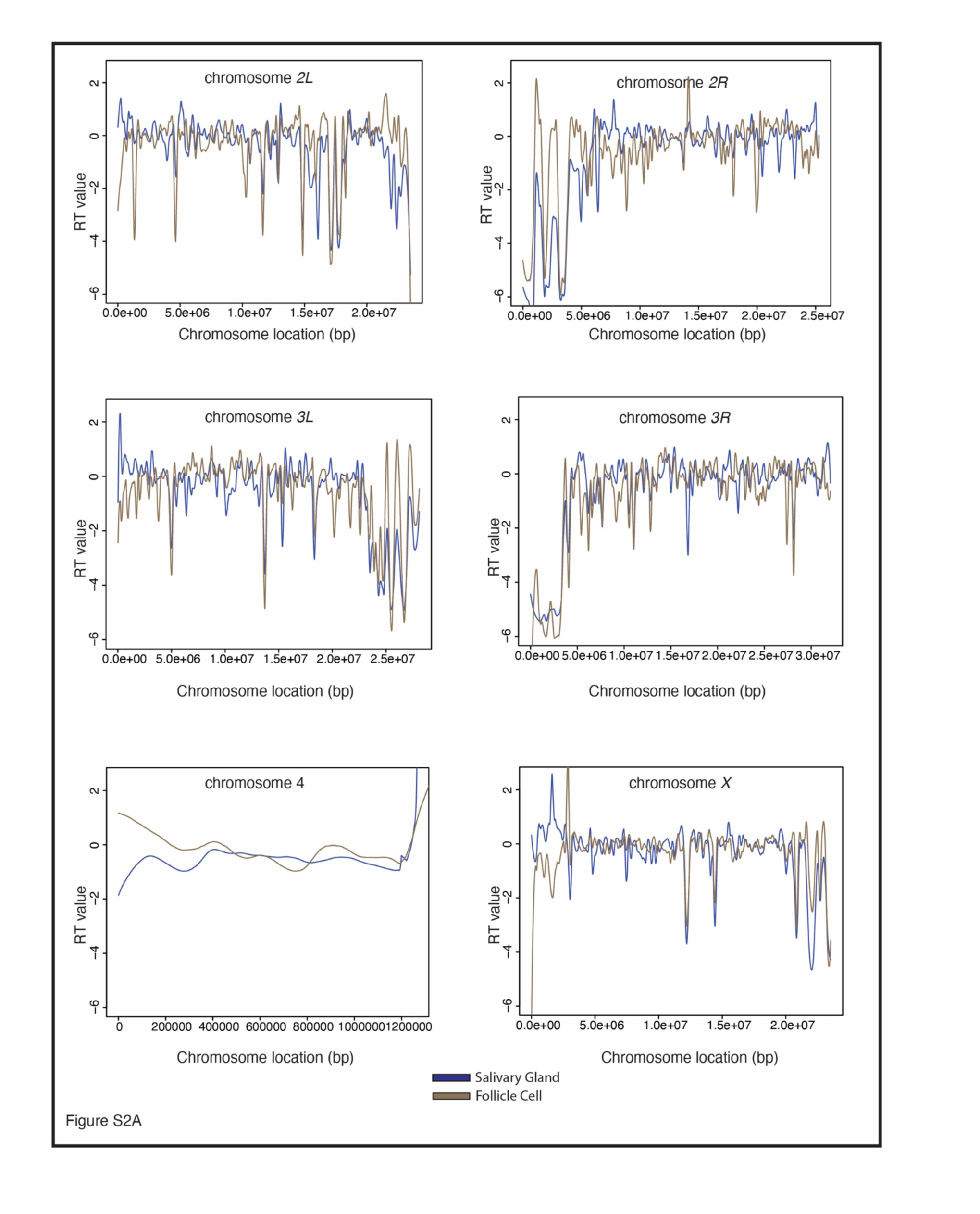
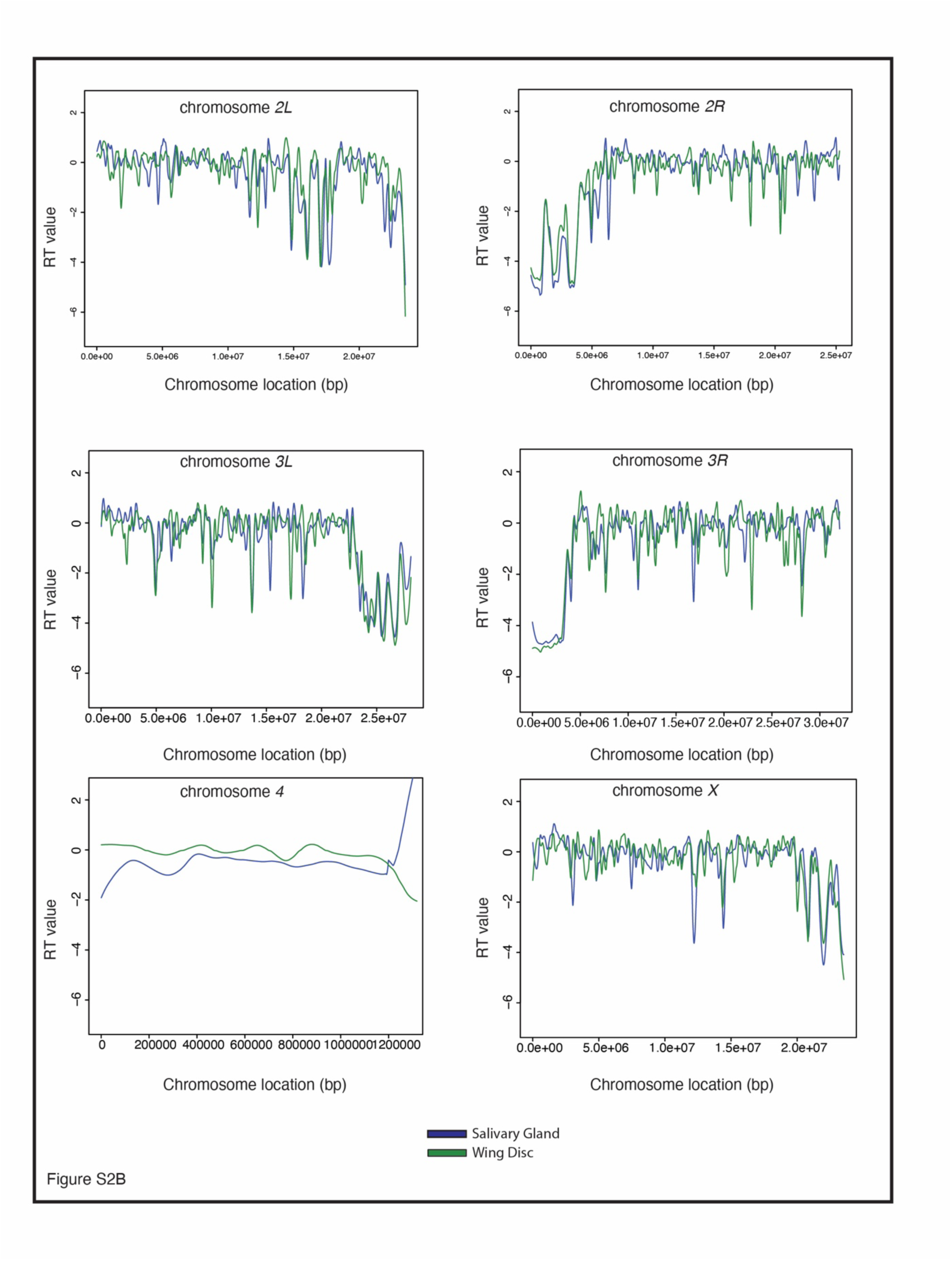
RT profiles generated with TIGER correlate with RT profiles generated by the G1/S method. A. LOESS regression lines showing RT values for wild-type salivary gland {blue) and wild-type follicle cell {brown)­ across all major chromosome scaffold. Each line represents the average of two biological replicates. B. LOESS regression lines showing RT values for wild-type salivary gland {blue) and wild-type wing disc (green)­ across all major chromosome scaffold. Each line represents the average of two biological replicates.

**Figure S3.**
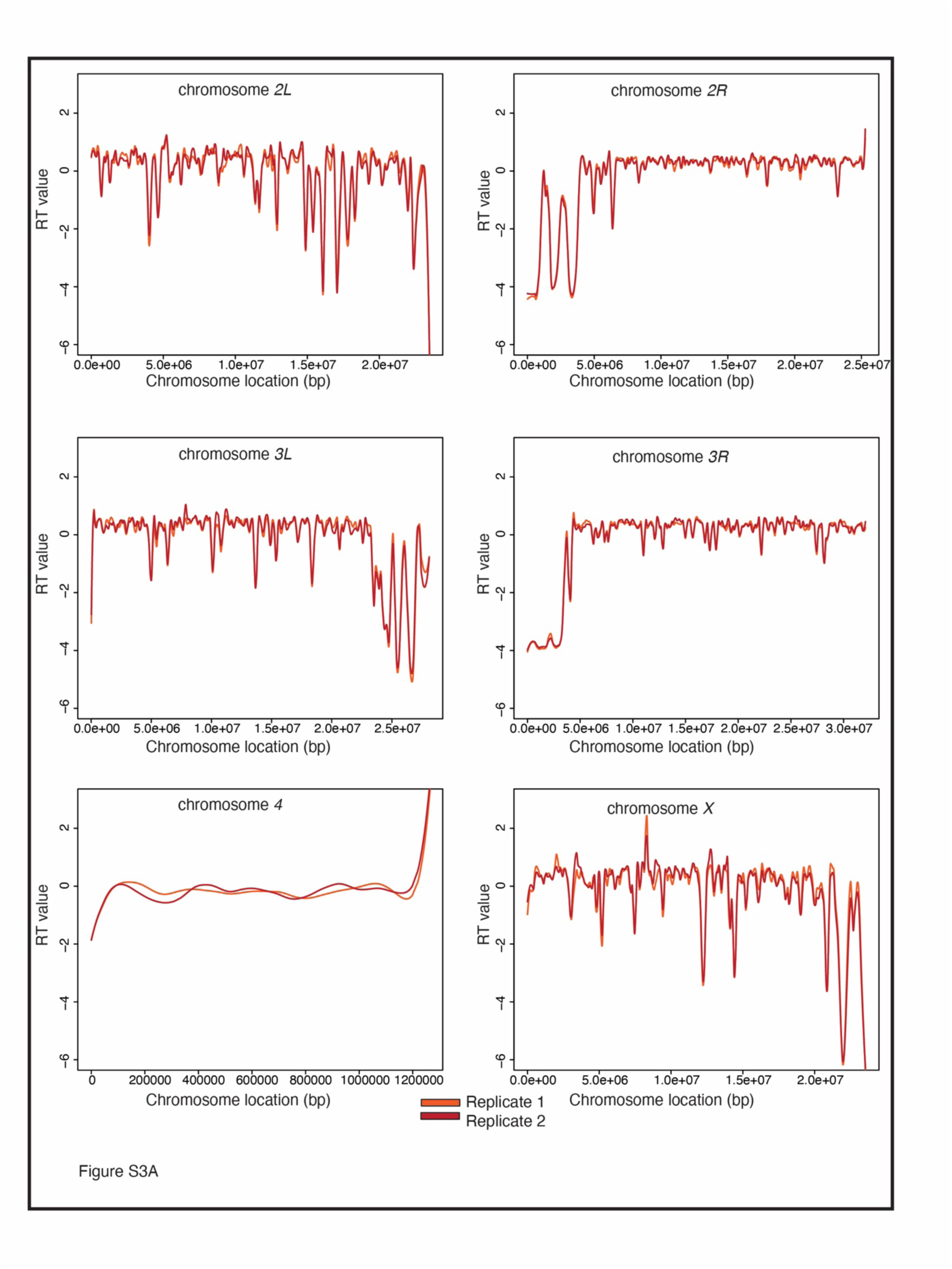
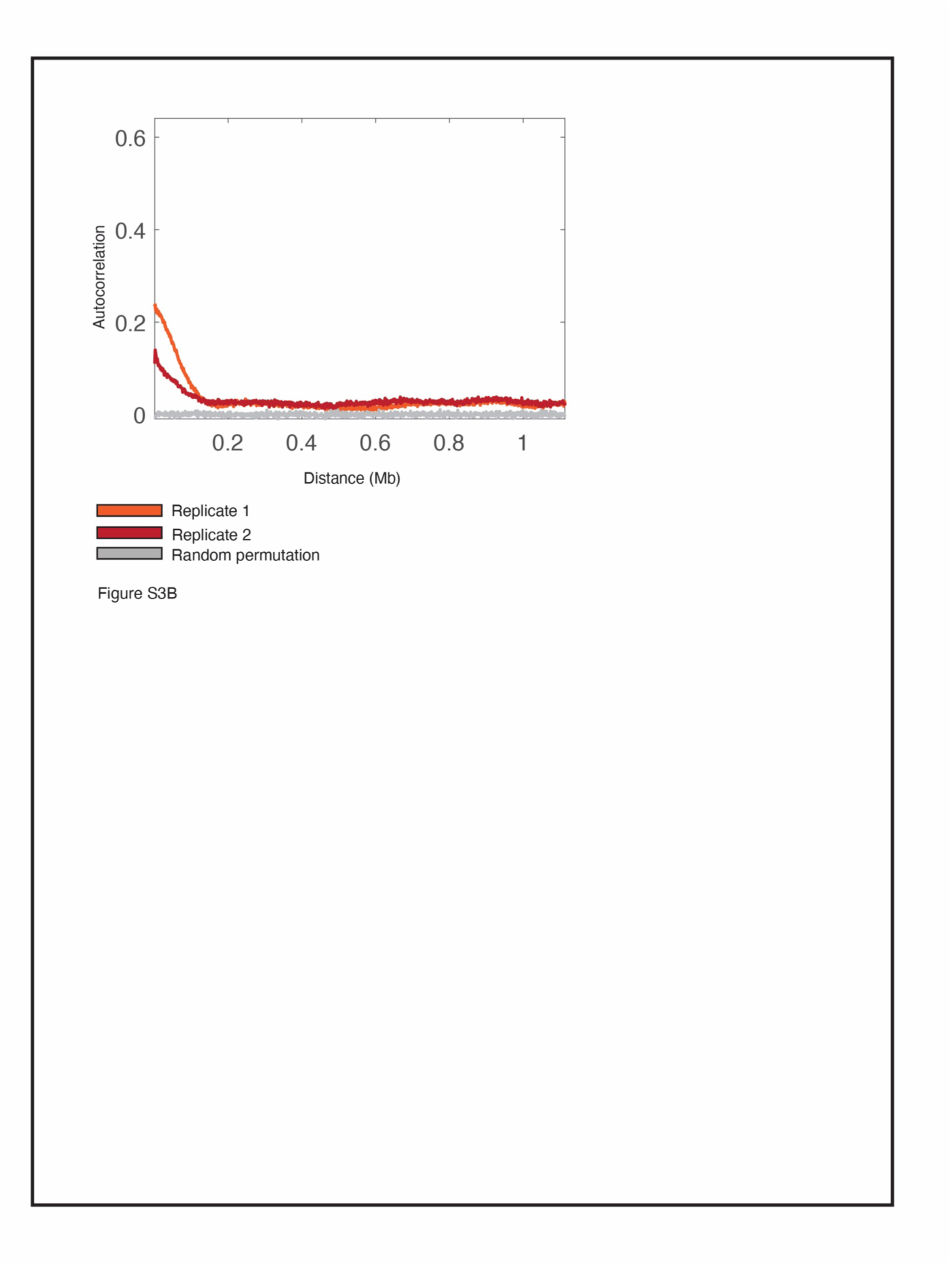
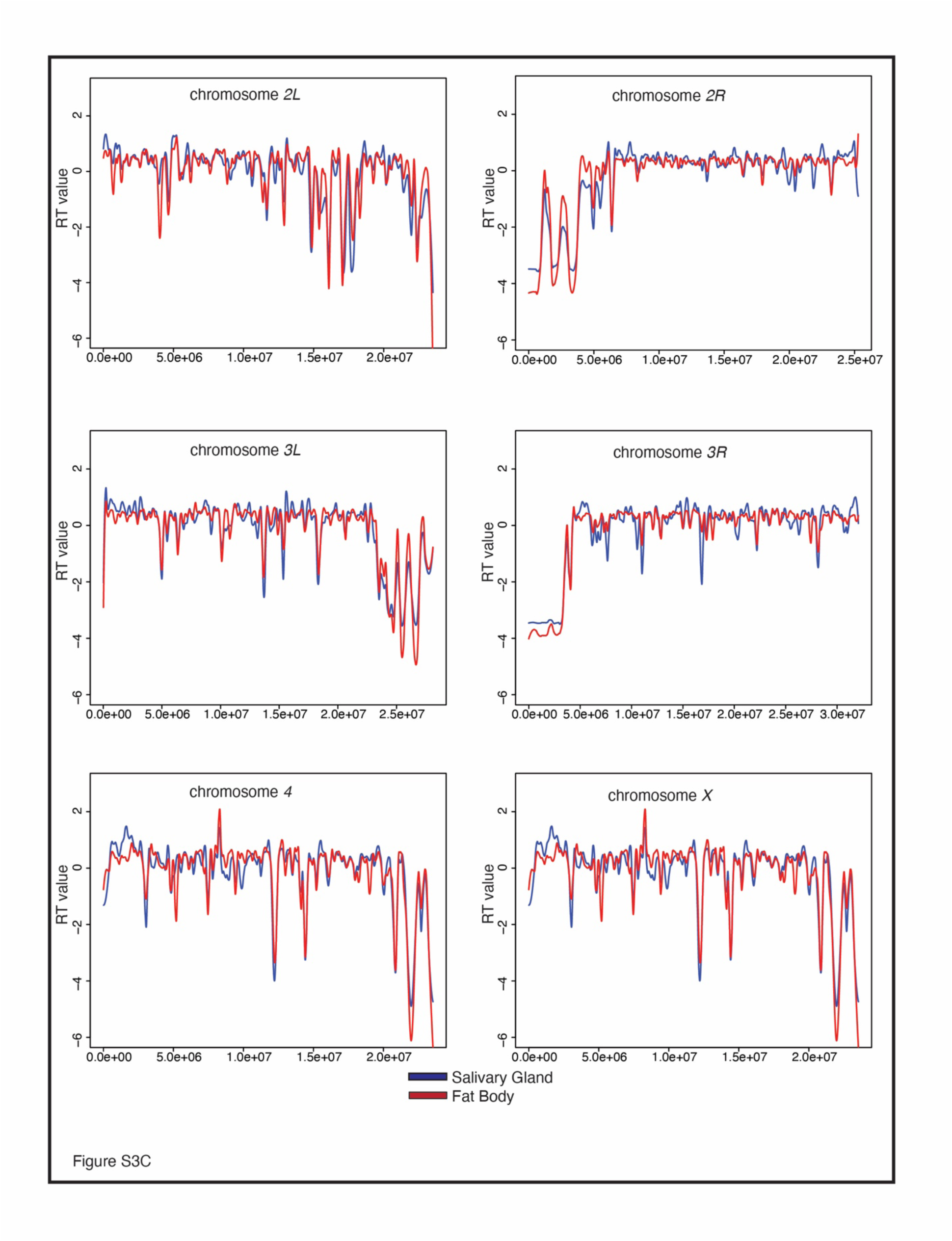
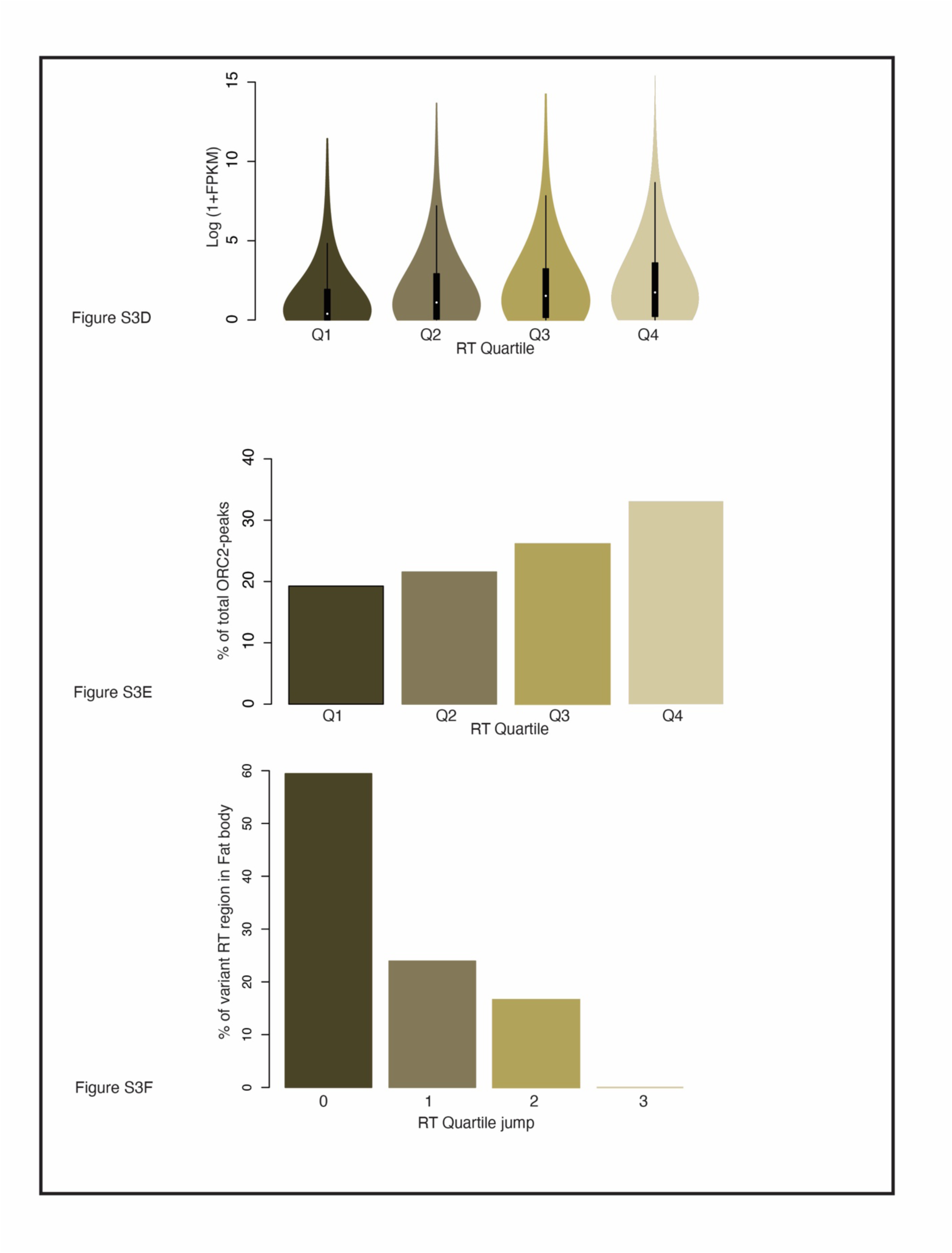
Tissue-specific RT correlates with tissue-specific underreplication. A. LOESS regression lines showing RT values for two biological replicates of wild-type fat bodies-replicate 1 (light red) and replicate 2 {dark red)- across all major chromosome scaffold. B. Autocorrelation values plotted for two replicates of wild-type fat body-replicate 1 (light red) and replicate 2 (dark red). Grey line depicts autocorrelation values for random permutation of RT values. C. LOESS regression lines showing RT values for wild-type salivary gland (blue) and wild-type fat body (red)- across all major chromosome scaffold. Each line represents the average of two biological replicates. D. RTwindows from fat body were divided into quartiles withthe lowest rawRT values in 01 and the highest rawRT values in Q4. Average transcript **FPKM** values were calculated for every transcript within RT windows. The log2-transformed (1+FPKM) values were plotted for transcripts in each RT quartile in a violin plot. E. Bar graph showing the % of ORC2-peaks corresponding to RT windows grouped into RT quartiles. F. Bar graph showing the % of variant RT regions that jumps either 0, 1, 2 or 3 quartiles between salivary gland and fat body.

**Figure S4.**
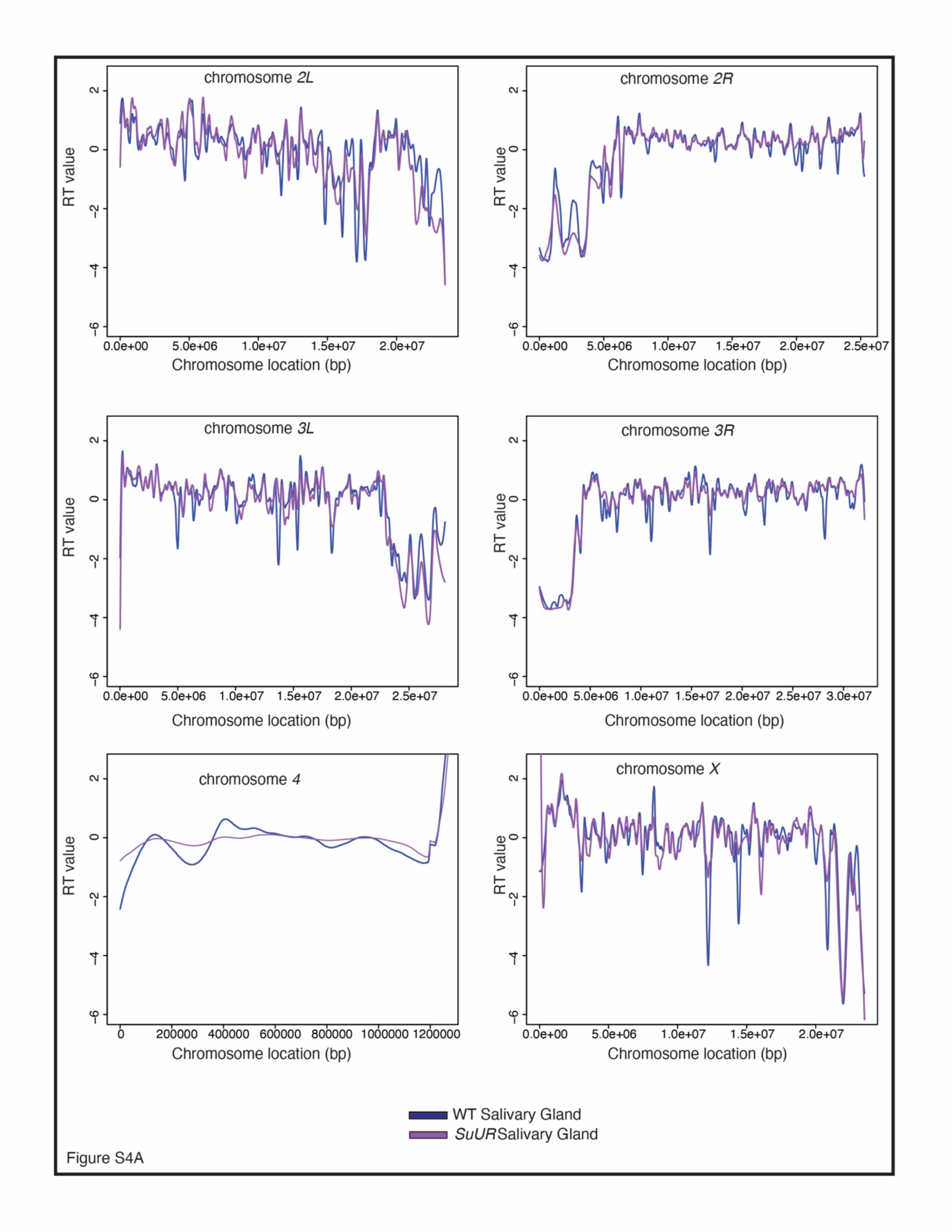
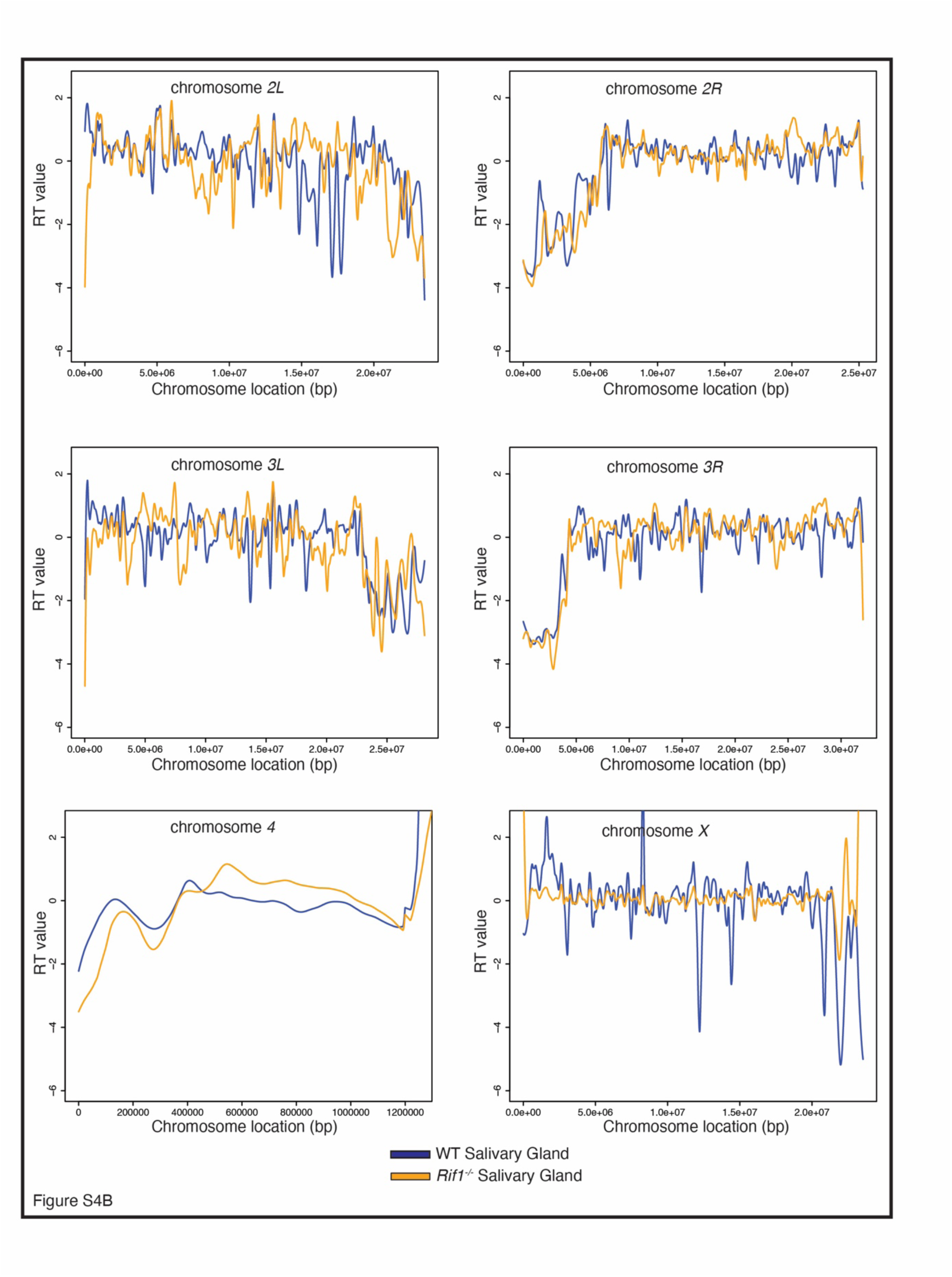
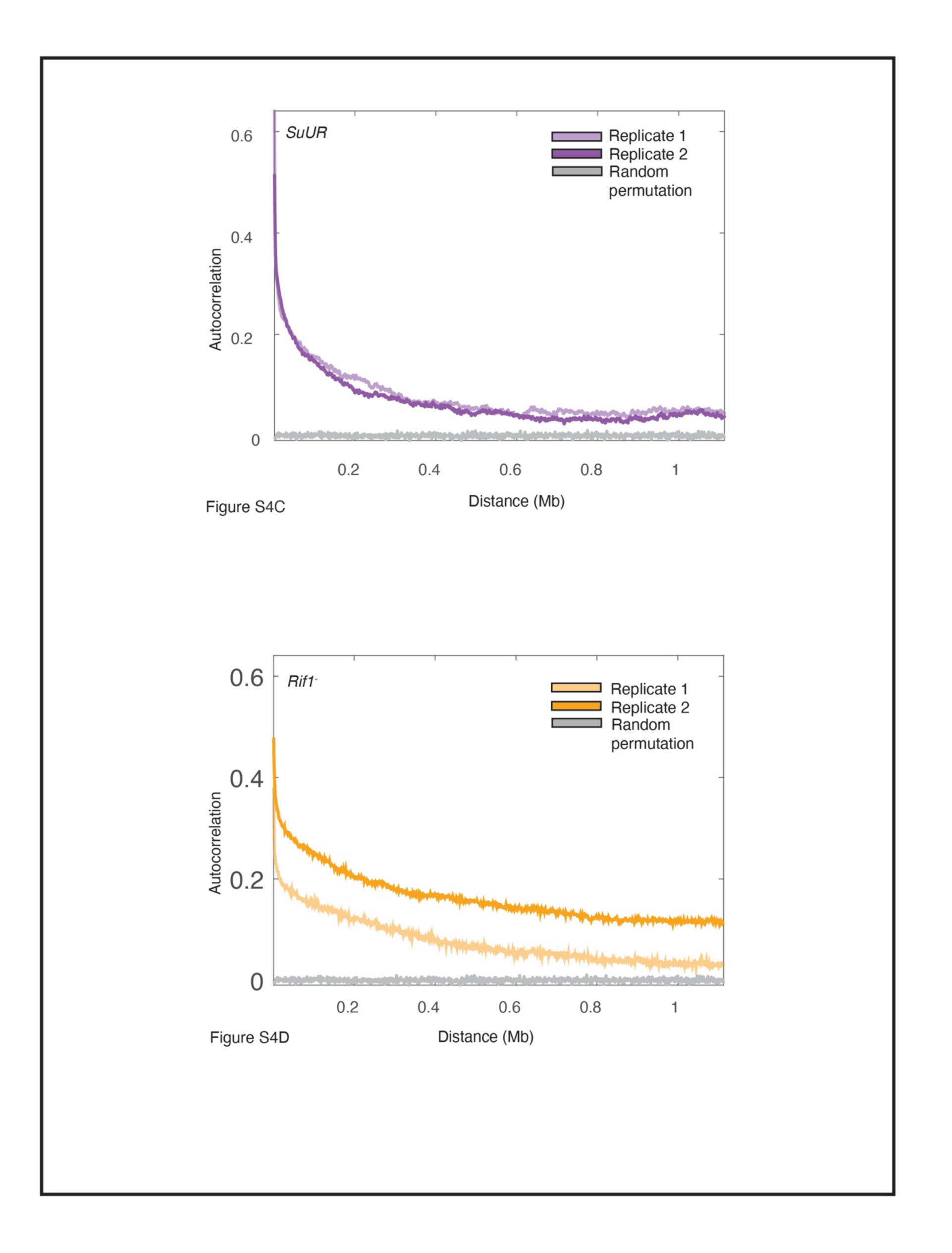
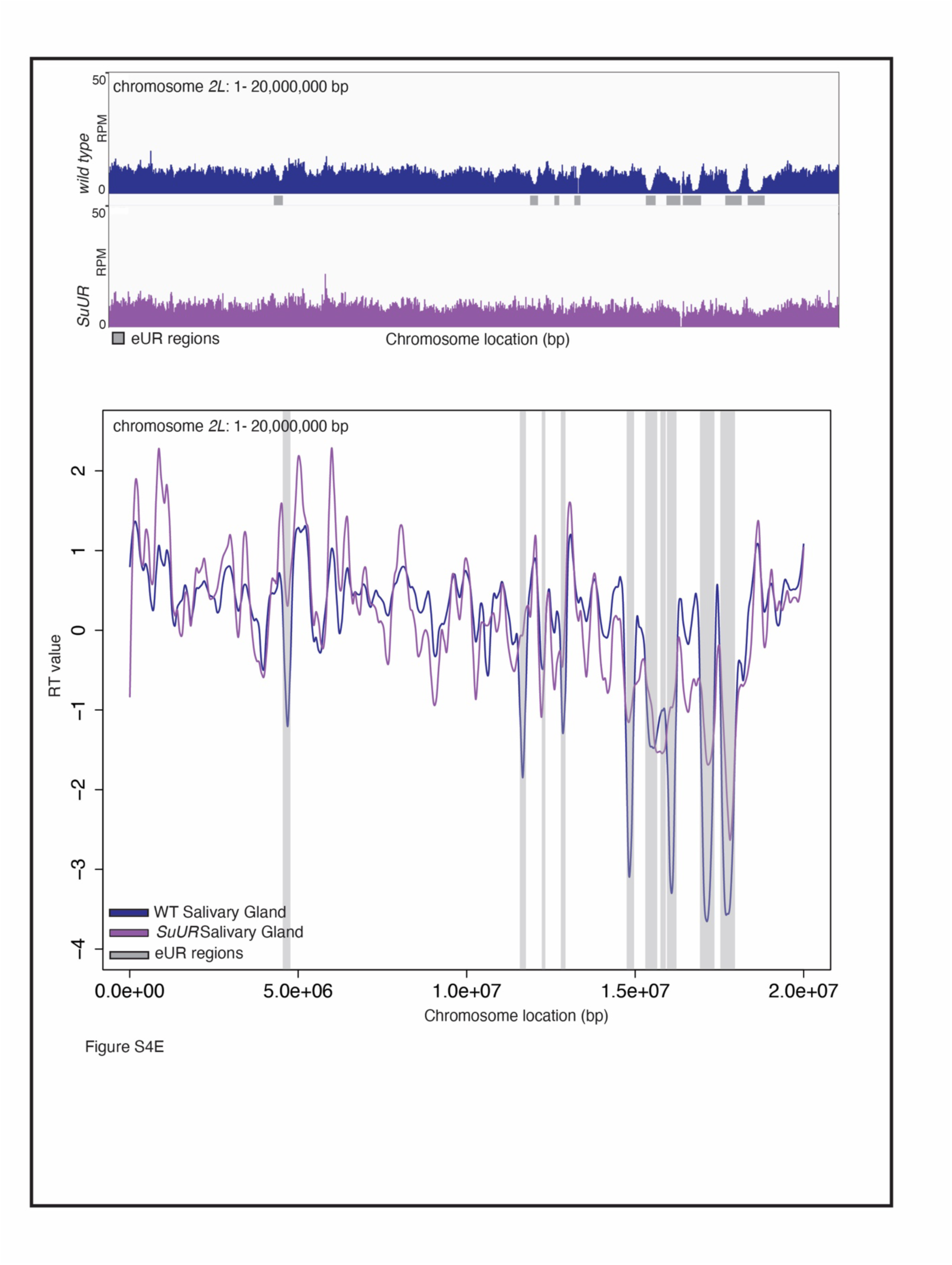
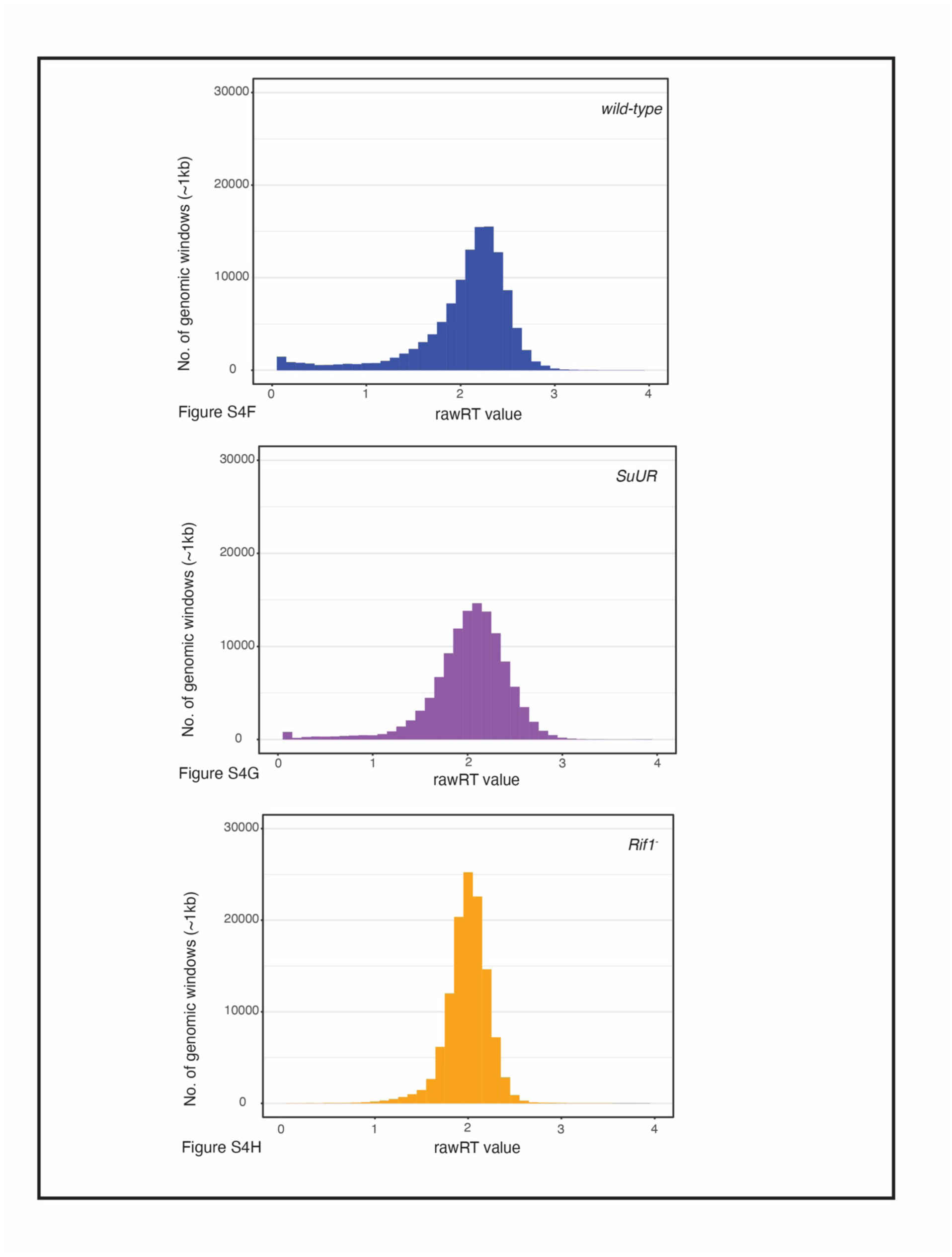
SUUR and Rif1 have differential effects on RTand UR. A. LOESS regression lines showing RT values for wild-type (blue) and *SuUR* mutant (purple) salivary gland, across all major chromosome scaffold. Eachline represents the average of two biological replicates. B. LOESS regress ion lines showing RTvalues for wild-type {blue) and *Rif1* mutant (yellow) salivary gland, across all major chromosome scaffold. Eachline represents the average of two biological replicates. C. Autocorrelation values plotted for two replicates of *SuUR* salivary gland-replicate 1 (light purple) and replicate 2 (dark purple). Grey line depicts autocorrelation values for random permutation of RT values. D. Autocorrelation values plotted for two replicatesof *Rif1* salivary gland-replicate 1(light yellow) and replicate 2 {dark yellow). Grey line depicts autocorrelationvalues for random permutation of RT values. E. lllumina-basedcopy number profiles (Reads Per Million; RPM) of chr2L:1 - 20,000,000 base pairs from wild type and *SuUR* mutant larval salivary glands. Grey bars below each profile represent euchromatic underreplicatedregions (eUR). F. Histogram quantifying the distribution of rawRT values in wild-type (blue) salivary gland. Bin width = 0.1. G. Histogram quantifying the distribution of rawRT values in *SuUR* (purple) mutant salivary gland. Bin width= 0.1. H. Histogram quantifying the distribution of rawRT values in *Rif1* (yellow) mutant salivary gland. Bin width = 0.1.

## Notes

https://www.ncbi.nlm.nih.gov/geo/query/acc.cgi?acc=GSE172375

### Competing Interest Statement

The authors have declared no competing interest.

